# Blastocoel expansion and AMOT degradation cooperatively promote YAP nuclear localization during epiblast formation

**DOI:** 10.1101/2024.08.15.608017

**Authors:** Hinako Maeda, Hiroshi Sasaki

## Abstract

The epiblast is a pluripotent cell population formed in the late blastocyst stage of preimplantation embryos. During the process of epiblast formation from the inner cell mass (ICM) of the early blastocyst, activation of the Hippo pathway transcription factor TEAD by the nuclear translocation of the coactivator protein YAP is required for the robust expression of pluripotency factors. However, the mechanisms that alter YAP localization during epiblast formation remain unknown. Here, we reveal two such mechanisms. Expansion of the blastocoel promotes nuclear YAP localization by increasing cytoplasmic F-actin and reducing YAP phosphorylation. Additionally, cell differentiation regulates YAP. Expression of the junctional Hippo component, AMOT, gradually decreases during epiblast formation through a tankyrase-mediated degradation. SOX2 expression in the ICM is necessary for the reduction of AMOT and YAP phosphorylation. These two mechanisms function in parallel. Thus, the blastocoel–F-actin and SOX2–AMOT axes cooperatively suppress YAP phosphorylation and promote YAP nuclear localization during epiblast formation. The cooperation of these two distinct mechanisms likely contributes to the robustness of epiblast cell differentiation.

## Introduction

During preimplantation development, mouse embryos form a cyst-like structure known as the blastocyst. The blastocyst consists of three types of cells: epiblast, primitive endoderm, and trophectoderm. The epiblast is a pluripotent tissue that later gives rise to the entire embryo and germ cells. The trophectoderm and primitive endoderm later form extraembryonic tissues, the placenta, and the yolk sac, respectively, supporting embryonic development (Cockburn and Rossant, 2010; Rossant and Tam, 2009). The epiblast is formed through two rounds of cell differentiation. The first differentiation occurs from the morula to the early blastocyst stage, during which the totipotent blastomeres differentiate into the inner cell mass (ICM) and the trophectoderm. Then, the ICM cells further differentiate into the epiblast and the primitive endoderm during the blastocyst stage (Cockburn and Rossant, 2010; Rossant and Tam, 2009). The Hippo signaling pathway plays important roles in both differentiation processes (Hashimoto and Sasaki, 2019; Nishioka et al., 2009).

The Hippo pathway is a conserved signaling pathway that regulates gene expression through downstream coactivator proteins YAP1 and TAZ/WWTR1 (hereafter collectively referred to as YAP) and TEA domain (TEAD) transcription factors TEAD1/2/3/4 (Chan et al., 2009; Lei et al., 2008; Ota and Sasaki, 2008; Zhao et al., 2007; Zhao et al., 2008). The Hippo pathway is regulated by various upstream signals, including molecular signals and mechano-physical status, such as cell-to-cell adhesion, cell-to-matrix adhesion, cell polarity, GPCR signaling, and mechanical forces that regulate the cytoskeleton (reviewed in (Ma et al., 2019; Zheng and Pan, 2019). Most of the upstream signals regulate the subcellular localization of YAP through phosphorylation by protein kinases LATS1/2, whereas mechanical cues also directly regulate YAP independently of LATS1/2 (Driscoll et al., 2015; Dupont et al., 2011; Elosegui-Artola et al., 2017).

In the first cell differentiation in preimplantation embryos, Hippo signaling regulates cell fates. Cell polarity, cell-cell adhesion, and mechano-physical status regulate Hippo signaling (Anani et al., 2014; Hirate et al., 2013; Maitre et al., 2016; Nishioka et al., 2009; Skory et al., 2023). The Hippo-inactive cells, which are located at the outer position in the embryo, accumulate the coactivator protein YAP in the nuclei, and TEAD4–YAP promotes trophectoderm differentiation by inducing trophectoderm-specific transcription factors, CDX2 and GATA3 (Nishioka et al., 2009; Ralston et al., 2010). The Hippo-active cells located at the inner position exclude YAP from the nuclei (Nishioka et al., 2009). The absence of nuclear YAP allows the expression of a pluripotency transcription factor, SOX2, via de-repression, promoting ICM fate (Frum et al., 2019; Nishioka et al., 2009). In the second differentiation, or the differentiation of the epiblast from the ICM, TEAD–YAP is involved in the expression of pluripotency factors (Hashimoto and Sasaki, 2019). In the ICM of the early blastocyst stage, YAP is in the cytoplasm. Then, YAP gradually accumulates in the nuclei, indicating the gradual increase of TEAD–YAP activity during epiblast formation between the mid and late blastocyst stages. The activation of TEAD–YAP is required for the strong expression of pluripotency factors in the epiblast cells (Hashimoto and Sasaki, 2019). Thus, the dynamic changes in YAP localization are important for proper epiblast development. However, the mechanisms that promote the nuclear localization of YAP during epiblast formation remain elusive.

In this study, we identified the involvement of two mechanisms. Mechano-physical signals of blastocoel expansion promote nuclear YAP, mediated by the increase of cytoplasmic F-actin and reduction of YAP phosphorylation. YAP is also regulated by cell differentiation. Expression of SOX2 in the ICM downregulates AMOT, leading to the reduction of YAP phosphorylation. These two mechanisms act in parallel and cooperatively suppress YAP phosphorylation to promote YAP nuclear localization during epiblast formation. The cooperation of these two distinct mechanisms likely contributes to the robustness of epiblast cell differentiation.

## Materials and Methods

### Mice

B6D2F1/Slc (referred to as BDF1) mice were either purchased from Japan SLC (Hamamatsu, Japan) or maintained in-house at the Animal Facility of the Frontier of Biosciences, University of Osaka. All experiments involving mice and recombinant DNA were conducted in accordance with the guidelines and approved protocols from the Animal Care and Use Committee of the Graduate School of Frontier Biosciences, University of Osaka, and the Gene Modification Experiments Safety Committee of the University of Osaka.

### Embryo collection and culture

Preimplantation embryos were collected following a standard protocol with slight modifications (Behringer et al., 2014). B6D2F1/Slc female mice were superovulated by intraperitoneal injection of 10 IU pregnant mare serum gonadotropin (ASKA Animal Health) and human chorionic serotropin (ASKA Animal Health) at 48-hour intervals, and then crossed with B6D2F1/Slc male mice to obtain wild-type embryos. One-cell-stage zygotes were collected from the E0.5 oviduct ampulla and treated with hyaluronidase (Sigma-Aldrich, H4272) to remove cumulus cells. Two-cell-stage embryos were subsequently collected from the E1.5 oviduct. All embryos were cultured in 10 μL drops of KSOM medium (ARK-Resource, I0BAIK200) covered with 5 μL of mineral oil (Sigma-Aldrich, M8410) in each well of a 72-well Nunc MiniTrays with Nunclon Delta Surface (Nunc 136528), maintained at 37 °C in a 5% CO2 incubator.

### sgRNA Design and synthesis

The sgRNAs targeting *Lmna* and *Sox2* genes were designed using CHOPCHOP (Labun et al., 2019). Template DNA preparation, in vitro transcription, and sgRNA purification followed previously described methods (Hashimoto et al., 2016). Briefly, template DNA fragments for each sgRNA were generated in separate PCR reactions using forward primers specific to each target gene, a common reverse primer, and the pX330-U6-Chimeric_BB-CBh-hSpCas9 plasmid DNA (Addgene, 42230) as template (Cong et al., 2013). KOD Fx neo DNA polymerase (KFX-201, Toyobo, Osaka, Japan) was employed under the following PCR conditions: 96°C for 1 min, followed by 40 cycles of 96°C for 10 s, 63°C for 10 s, 68°C for 30 s, and final extension at 68°C for 1 min. Template DNA was purified by ethanol precipitation and used for sgRNA synthesis with the MEGAshortscript T7 Transcription Kit (AM1354, Thermo Fisher Scientific). sgRNAs targeting the same gene were pooled, purified by phenol extraction and isopropanol precipitation, and dissolved in RNase-free water. The following primers (Eurofins Genomics) were used for PCR reactions: *DsRed* gRNA Forward-1: 5’-TTAATACGACTCACTATAGGGGCCACGAGTTCGAGATCGAGTTTTAGAGCTAGAAA-3’

*Lmna* gRNA Forward-1: 5’-TTAATACGACTCACTATAGGCGAGCTCCATGACCTGCGGTTTTAGAGCTAGAAATAGCAAGTTAAAAT-3’

*Lmna* gRNA Forward-2: 5’-TTAATACGACTCACTATAGGGGGACTTGTTGGCTGCGCGTTTTAGAGCTAGAAATAGCAAGTTAAAAT-3’

*Sox2* gRNA Forward-1: 5’-TTAATACGACTCACTATAGGGCCCGCAGCAAGCTTCGGGTTTTAGAGCTAGAAATA

GCAAGTTAAAAT-3’

*Sox2* gRNA Forward-2: 5’-TTAATACGACTCACTATAGGGCCTCAACGCTCACGGCGGTTTTAGAGCTAGAAATAGCAAGTTAAAAT-3’

*Sox2* gRNA Forward-3: 5’-TTAATACGACTCACTATAGGGCTCTGTGGTCAAGTCCGGTTTTAGAGCTAGAAATAGCAAGTTAAAAT-3’

Common gRNA Reverse: 5’-AAAAGCACCGACTCGGTGCCACTTTT-3’

### Genome editing of zygotes

Genome editing of zygotes was conducted by introducing Cas9 protein (Integrated DNA Technologies) along with the synthesized sgRNAs using electroporation, following established procedures (Hashimoto et al., 2016).

### Fluorinert injection into blastocoel

Fluorinert FC-40 (Sigma, F9755-100ML) was injected into the blastocoel of early-mid blastocyst stage embryos using an injection needle and a micromanipulator system. The injection procedure applied pressure to the inner cell mass while ensuring the trophectoderm and the zona pellucida remained intact. Subsequently, the injected embryos were cultured in KSOM medium for 15 minutes before being fixed in 4% paraformaldehyde (PFA) in phosphate-buffered saline (PBS) for another 15 minutes at room temperature. Following fixation, the injected Fluorinert was removed by creating a hole in the mural trophectoderm to facilitate observation of immunofluorescent signals.

### Blastocoel suction

Blastocoel fluid from late-mid blastocyst stage embryos was suctioned using an injection capillary with a diameter of approximately 5 μm, employing a micromanipulator system. Once the compression of the blastocoel was confirmed, the embryos were cultured in KSOM medium for 15 minutes. Subsequently, they were fixed in 4% PFA in PBS for 15 minutes at room temperature.

### Immunofluorescent staining and confocal image acquisition

Immunofluorescent staining of embryos was performed as described previously (Hashimoto and Sasaki, 2019). All procedures were performed at room temperature unless otherwise noted. Briefly, embryos were fixed in 4% PFA in PBS for 15 minutes, washed, and permeabilized with 0.1% Triton X-100 in PBS (PBST) for 1 minute, twice. The embryos were then blocked with 2% donkey serum in PBST (blocking solution) and incubated overnight with primary antibodies diluted at 1:100 in the blocking solution. For detection of p-YAP signals, embryos were fixed with ice-cold methanol for 20 minutes on ice. Phosphatase Inhibitor Cocktail (Nacalai Tesque, 07574-61) was added at a 100-fold dilution from the post-fixation washing step to the primary antibody reaction. The primary antibodies used are as follows: rabbit monoclonal anti-Yap1 antibody (1:200) (Cell Signaling Technology, D8H1X), rabbit monoclonal anti-Sox2 antibody (1:200) (Cell Signaling, D9B8N), rabbit polyclonal anti-Amot antibody (Amot-C, 1:200) (Hirate et al., 2013), rabbit polyclonal anti-p-Yap (Ser127) antibody (1:200) (Cell Signaling, 4911), rat monoclonal anti-E-cadherin antibody (ECCD-2) (1:200) (Takara, Cat# M108), mouse monoclonal anti-Yap1 antibody (MO1) (1:200) (Abnova, H00010413-M01), mouse monoclonal anti-Lamin A/C antibody (1:100) (BD Transduction Laboratories, 612162), goat polyclonal anti-Sox2 (R&D, AF2018). After washing in PBST for 1 minute, twice, the embryos were incubated with the secondary antibodies and Hoechst 33342 at a 1:1000 dilution in PBST for more than 2 hours. The secondary antibodies used are as follows: donkey anti-mouse IgG 488/555 (Invitrogen, A32766/A32773), donkey anti-rabbit IgG 488/555/647 (Invitrogen, A32790/A32794/A21245), donkey anti-goat IgG 488/647 (Invitrogen, A-11055/A21447), donkey anti-rat IgG 488 (Invitrogen, A21208), Alexa Fluor 647 phalloidin (1:50) (Molecular Probes, A22287). Immunofluorescence-stained embryos were placed in a drop of PBS on a glass base dish, and confocal images were obtained using a Nikon A1 point-scanning inverted confocal microscope (Nikon Solutions Inc., Tokyo, Japan) or a spinning disk confocal microscope system consisting of Nikon Ti2 inverted microscope (Nikon Solutions Inc., Tokyo, Japan), CSU-W1 confocal unit (Yokogawa Electric Corp., Tokyo, Japan), and an ORCA-Flash4.0 sCMOS camera (Hamamatsu Photonics K.K., Shizuoka, Japan). Super-resolution images were acquired using the LSM900 (ZEISS) microscope equipped with Airyscan1 and analyzed with the ZEN lite software (ZEISS).

### Image analysis

Confocal images were analyzed with NIS-Elements AR analysis (Nikon Solutions, Tokyo, Japan) or Imaris (Bitplane, Belfast, UK) software.

### Cell number measurement

Cell number was counted using Imaris software. A 3D image was constructed from a series of confocal images. The number of nuclei in an embryo stained with Hoechst was counted using the spot function of the object tool. Dividing cells were counted as a single cell, and apoptotic cells were excluded.

### Measurement of fluorescent signals

Z-stack confocal images acquired with Z-intervals of 1 µm were analyzed using Imaris software. Signal intensities were obtained by manual segmentation and 3D reconstruction of the regions of interest (ROIs) in the target area using the surfacing function of the object tool. Four different types of segmentations were performed based on the signals to be acquired. To measure the nuclear signals (e.g., nuclear YAP signals), the nuclear area of the SOX2-positive ICM/epiblast cells was segmented (Fig. S6A). The average YAP signal intensity was calculated for each ROI. Attenuation of signal intensities along the Z-axis was compensated by dividing the nuclear signals by the corresponding Hoechst signals. To measure the cytoplasmic signals (e.g., cytoplasmic F-actin), the cytoplasmic area between the strong junctional F-actin signal and the nuclear Hoechst signal was manually segmented as ROIs. ROIs were set to avoid overlap with the junctional F-actin and the Hoechst signals (Fig. S6B). Attenuation of signal intensities along the Z-axis was compensated by dividing the cytoplasmic signals by the corresponding cells’ Hoechst signals. To measure the p-YAP and AMOT signals, entire ICM/epiblast regions marked by SOX2 expression were manually segmented as ROIs. E-cadherin signals were used to identify the boundaries of the cells. For p-YAP signals, segmentation boundaries were set inside the outermost E-cadherin signals (Fig. S6C). For AMOT signals, segmentation boundaries were set on the outermost E-cadherin signals (Fig. S6D). The average signal intensities for p-YAP and AMOT were calculated for each ROI. Attenuation of signal intensities along the Z-axis was compensated by dividing p-YAP/AMOT signals by the corresponding averaged Hoechst signals.

### Measurement of the nuclear sphericity

Sphericity is the ratio of the surface area of a sphere (with the same volume as the particle) to the surface area of the particle itself (Wadell, 1933). The sphericity values of the SOX2-positive nuclei were calculated using the 3D surface model produced by manual segmentation described above.

### Measurement of the blastocoel volume

Blastocyst volume was measured using Imaris software. Surface rendering of a blastocyst was first performed using E-cadherin signals. Weak background-level signals were also used to identify the entire cell area. This procedure generated a surface model of a blastocyst enclosing all the cells and excluding the blastocoel. Voxels outside the surface model were selected and used to create a new channel. Surface rendering of the new channel generated new surface models outlining the voxels outside the cells. The inner surface model corresponded to the blastocoel. The outer surface model was ignored. The volume of the inner surface model was measured as the blastocoel volume.

### XAV939 treatment

Early-mid blastocysts were cultured in KSOM medium containing 10 μM XAV939 (Sigma-Aldrich, X3004) for 12 hours. Control embryos were cultured in KSOM medium containing the same concentration of dimethyl sulfoxide (DMSO).

### Statistical Analysis

Statistical analyses were performed using GraphPad Prism 9.0 software (GraphPad, La Jolla, CA, USA). The number of cells (n) or embryos (N) analyzed, error bars, statistical analyses, and *p*-values are all stated in each figure or figure legend.

## Results

### Blastocyst staging based on subcellular distribution of YAP in the ICM

To investigate the mechanisms of nuclear translocation of YAP during the blastocyst stage, we first defined the substages of blastocyst development based on the subcellular localization of YAP. YAP localization within the SOX2-positive ICM/epiblast cells was classified into the following three categories (Fig. 1A): cytoplasm, where the YAP signal is predominantly in the cytoplasm; uniform, where the YAP signal is distributed evenly between the cytoplasm and nucleus; nucleus, where the YAP signal is predominantly in the nucleus. We assessed the distribution of these categories across the blastocyst stages and documented the proportions within the ICM/epiblast (Fig. 1B, C). Observations revealed that in embryos ranging from 33 to 68 cells, more than 80% of the cells exhibited a cytoplasmic pattern. Between 72 and 97 cells, YAP localization was highly variable. The proportion of cytoplasmic cells gradually decreased, and the ratio of nuclear cells increased as the total cell number increased. Beyond 102 cells, no cytoplasmic cells were observed. Based on these results, we defined the blastocyst stages as follows: early blastocyst up to 69 cells; mid blastocyst for 70 to 99 cells; and late blastocyst for over 100 cells. The mid blastocyst stage was further divided into early-mid (70 to 84 cells) and late-mid (85 to 99 cells) blastocyst stages (Fig. 1B, C).

**Figure 1.**
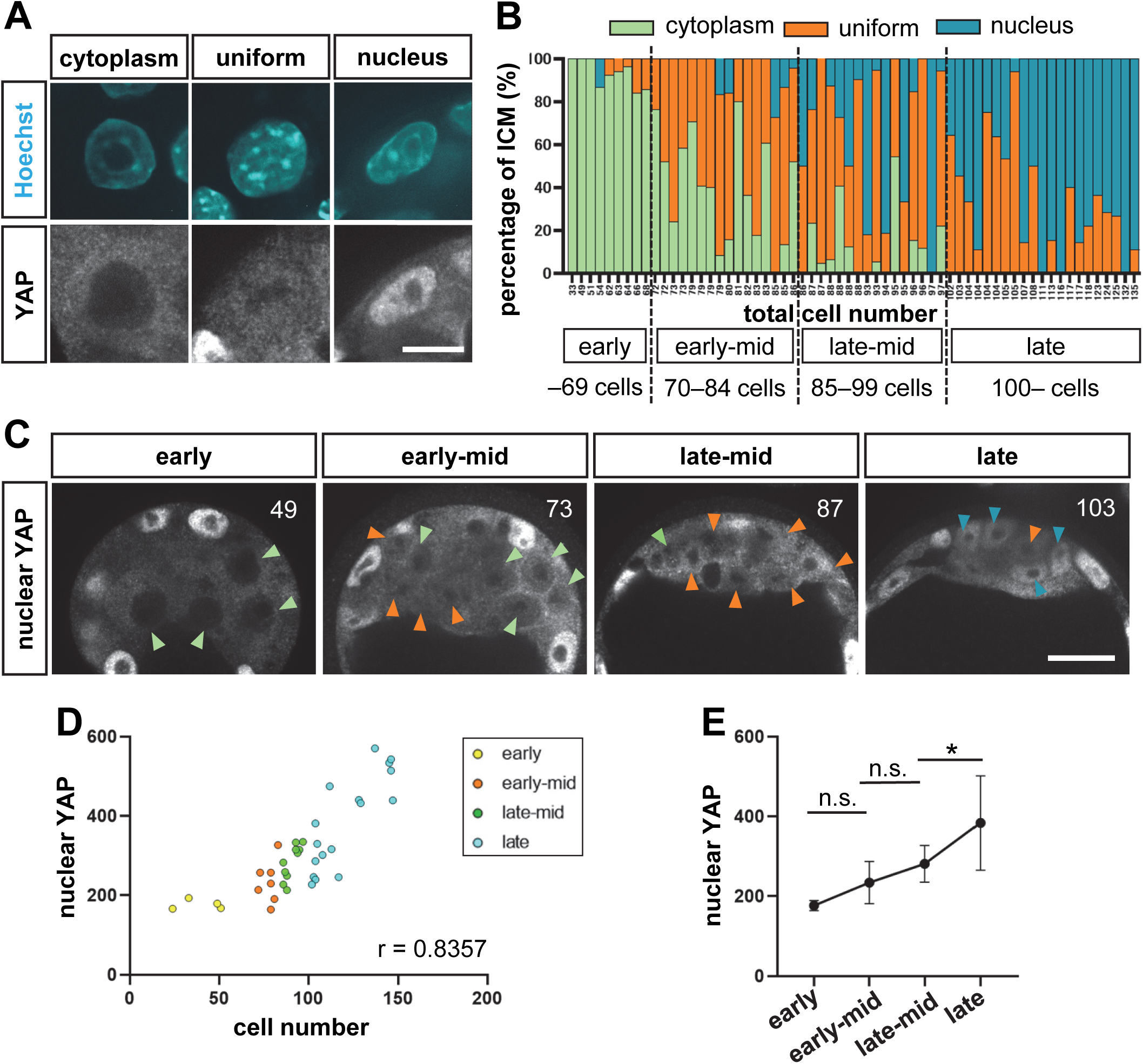
Blastocyst staging based on YAP distribution in the ICM/epiblast. (A) Representative immunofluorescence images showing different subcellular distributions of YAP. SOX2-positive ICM/epiblast cells were classified into three categories based on the subcellular distribution of YAP: cytoplasm (left), uniform (middle), and nucleus (right). Scale bar represents 10 μm. (B) Developmental changes in the composition of YAP distribution in ICM/epiblast cells. Green, orange, and blue represent the percentages of cells with cytoplasmic, uniform, and nuclear YAP distributions, respectively. (C) Representative immunofluorescence images of YAP distribution in the ICM region at different developmental stages. The total number of cells in the embryo is indicated in each image. Green, orange, and blue arrowheads indicate cells with cytoplasmic, uniform, and nuclear YAP distributions, respectively. Scale bar represents 20 μm. (D) A dot plot showing the correlation between total cell number and nuclear YAP signals in the ICM/epiblast. (E) A line graph showing changes in nuclear YAP signals during blastocyst stages. The values shown represent the means and standard errors. *p*-values were determined by one-way ANOVA, followed by Dunn’s multiple comparison test. \**p* < 0.05, n.s.: not significant. The sample numbers analyzed for each stage were as follows: early (N = 4, n = 59), early-mid (N = 7, n = 142), late-mid (N = 9, n = 174), and late (N = 17, n = 278) blastocyst.

Next, we quantified the YAP signal levels in the nuclei of SOX2-positive ICM/epiblast cells. For this purpose, we manually segmented nuclei and quantified nuclear signals in three dimensions (Fig. S5A). The results showed that the nuclear YAP signals gradually increased with the progression of development (Fig. 1D, E).

### Expansion of blastocoel contributes to the nuclear translocation of YAP within the ICM

To elucidate the mechanisms that promote the nuclear translocation of YAP, we first focused on the expansion of the blastocoel during the blastocyst stages. Mechanical signals are an upstream regulator of YAP (Dupont et al., 2011; Wada et al., 2011). Because blastocoel expansion increases cortical tension of the trophectoderm cells (Chan et al., 2019), we hypothesized that blastocoel expansion also applies mechanical forces to the ICM cells, promoting the nuclear localization of YAP. Measurement of the blastocoel volume confirmed a constant increase in the blastocoel volume during the blastocyst stages (Figs. 2A, B, S1A). Thus, blastocoel expansion occurs in parallel with the increase in nuclear YAP signals.

**Figure 2.**
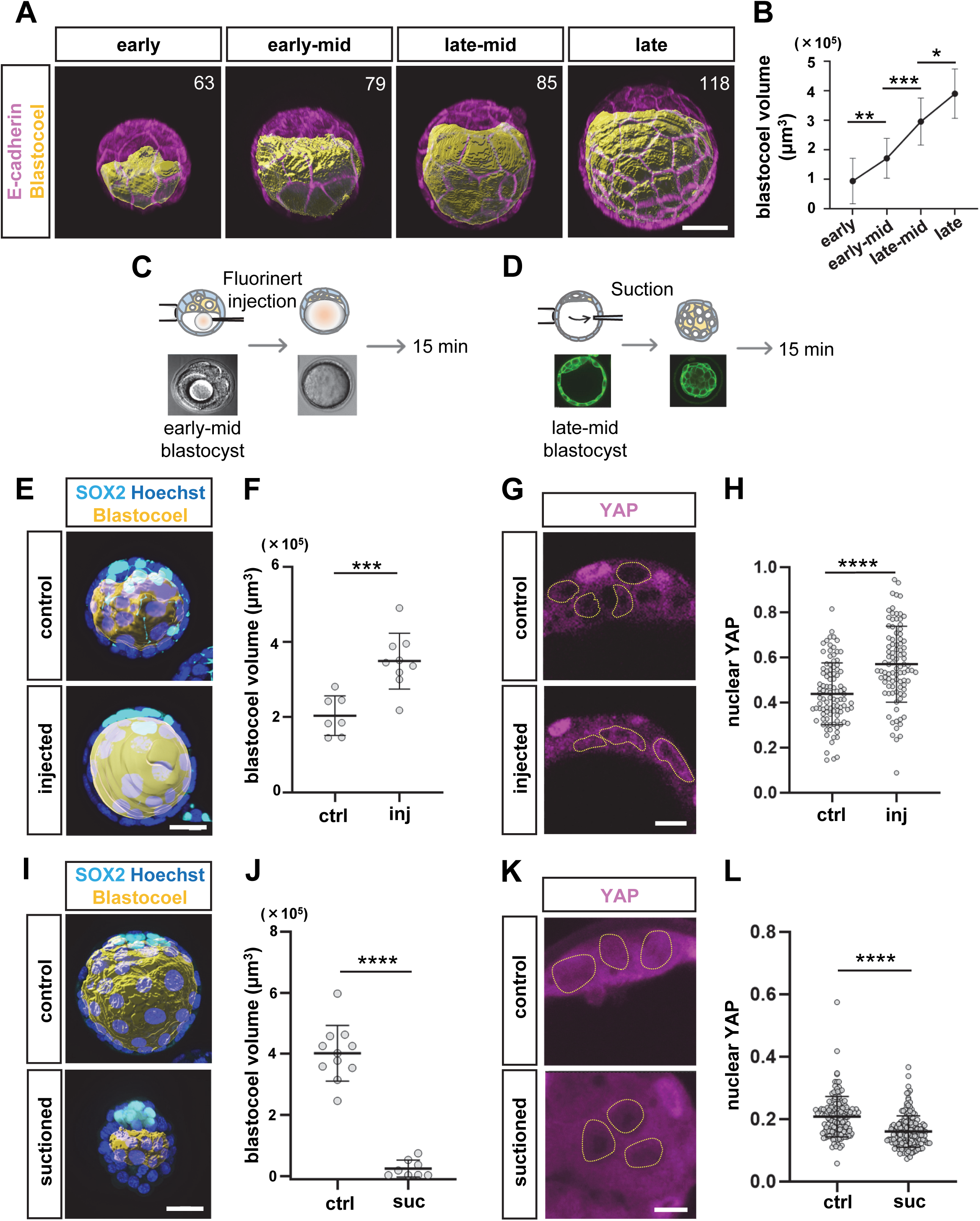
Blastocoel expansion promotes YAP nuclear localization. (A) Representative images of the results of surface rendering of blastocoel. The total number of cells in the embryo is indicated in the image. Orange and magenta represent blastocoel and E-cadherin signals, respectively. Scale bar represents 40 μm. (B) A line graph showing the changes in the blastocoel volume during blastocyst stages. The values shown represent the means and standard errors. *p*-values were determined by one-way ANOVA, followed by Dunn’s multiple comparison test. \**p* < 0.05, \*\**p* < 0.01, \*\*\**p* < 0.001. Sample numbers analyzed for each stage were as follows: early (N = 14), early-mid (N = 17), late-mid (N = 7), and late (N = 14) blastocyst. (C) Scheme of blastocoel expansion experiments. (D) Scheme of blastocoel shrinkage experiments. (E) Representative images of the blastocoel in blastocoel expansion experiments. Cyan, blue, and yellow represent SOX2, Hoechst, and blastocoel, respectively. Scale bar represents 20 μm. (F) Dot plots showing the changes in blastocoel volume with blastocoel expansion. Horizontal lines represent the means and standard errors. *p*-values were determined by Student’s *t*-test. \*\*\**p* < 0.001. Sample numbers analyzed for each group were as follows: control (N = 7), injected (N = 9). (G) Representative immunofluorescence images of YAP distribution in blastocoel expansion experiments. Dotted lines represent ICM nuclei. Scale bar represents 10 μm. (H) Dot plots showing the increase in nuclear YAP signals with blastocoel expansion. Horizontal lines represent the means and standard errors. *p*-values were determined by Student’s *t-*test. \*\*\*\**p* < 0.0001. Sample numbers analyzed for each group were as follows: control (n = 100), injected (n = 92). (I) Representative images of the blastocoel in blastocoel shrinkage experiments. Cyan, blue, and yellow represent SOX2, Hoechst, and blastocoel, respectively. Scale bar represents 20 μm. (J) Dot plots showing the changes in blastocoel volume with blastocoel suction. Horizontal lines represent the means and standard errors. *p*-values were determined by Student’s *t*-test. \*\*\*\**p* < 0.0001. Sample numbers analyzed for each group were as follows: control (N = 11), suctioned (N = 8). (K) Representative immunofluorescence images of YAP distribution in blastocoel shrinkage experiments. Dotted lines represent ICM nuclei. Scale bar represents 10 μm. (L) Dot plots showing the decrease in YAP N/C ratio with blastocoel suction. Horizontal lines represent the means and standard errors. *p*-values were determined by Student’s *t*-test. \*\*\*\**p* < 0.0001. Sample numbers analyzed for each group were as follows: control (n = 142), suctioned (n = 165).

To elucidate the relationship between blastocoel expansion and nuclear localization of YAP, we experimentally manipulated the blastocoel size. We first injected an inert liquid, Fluorinert, into the blastocoel of early-mid blastocysts to increase the blastocoel volume (Fig. 2C, E, F), and YAP distribution was examined 15 minutes after injection. We used Fluorinert to avoid leakage from the blastocoel. The nuclear YAP signal level in ICM cells was significantly increased in injected embryos compared to uninjected control embryos (Fig. 2G, H). To examine whether the observed increase in nuclear YAP signal in injected embryos is a consequence of nuclear translocation of YAP, we also assessed the nuclear to cytoplasmic (N/C) ratio of YAP signals. The N/C ratio was also increased, suggesting that blastocoel expansion increased nuclear YAP signals by promoting the translocation of YAP from the cytoplasm to the nucleus (Fig. S2A, B). As an opposite experiment, we next reduced the blastocoel volume by suctioning fluid from the blastocoel of late-mid blastocyst stage embryos (Fig. 2D, I, J). Fifteen minutes after suction, the nuclear YAP signal intensity of the suctioned embryos was significantly lower than that of control embryos (Fig. 2K, L), indicating that reduction of blastocoel volume excludes YAP from the nucleus. The blastocoel size reduction experiments suggest that the forces applied in normal (unmanipulated) embryos promote nuclear localization of YAP. Taken together, these results suggest that blastocoel expansion is a mechanism that promotes nuclear translocation of YAP during the mid blastocyst stage.

### Flattened nuclear morphology appears to have a minor role in the nuclear translocation of YAP

In cultured cells, a flattened nuclear morphology promotes the nuclear localization of YAP (Driscoll et al., 2015; Elosegui-Artola et al., 2017). To determine the importance of nuclear morphology on YAP distribution, we examined the sphericity of nuclei as a measure of nuclear morphology. A low sphericity indicates a flattened morphology. Nuclei were manually segmented to calculate the sphericity in three dimensions (Fig. S3A). The nuclear sphericity showed a tendency to decrease as the blastocyst stage progressed (Figs. S3B; S1B), suggesting an inverse correlation between nuclear sphericity and nuclear YAP overall. At the level of individual cells, however, the correlation between nuclear YAP signal intensity and nuclear sphericity was minimal at both early-mid (Fig. S3C) and late-mid (Fig. S3D) blastocyst stages. Therefore, nuclear morphology does not play a major role in the nuclear translocation of YAP in mid blastocyst stage embryos.

### Cytoplasmic F-actin is required for the nuclear localization of YAP

In cultured cells, cytoplasmic F-actin stress fibers play an important role in the regulation of YAP (Dupont et al., 2011; Wada et al., 2011; Zhao et al., 2012). Thus, we hypothesized that cytoplasmic F-actin regulates YAP during the mid blastocyst stage. To test this hypothesis, we first examined cytoplasmic F-actin signals of the SOX2-positive ICM/epiblast cells during blastocyst stages (Fig. 3A, B; S1C). Cytoplasmic F-actin was very low at the early blastocyst stage and gradually increased from early to late-mid blastocyst stages, coinciding with the timing of YAP nuclear translocation (Figs. 3A, B; S1C). Furthermore, F-actin signal intensity in individual cells correlated well with nuclear YAP signal intensity at both the early-mid (Fig. 3C) and late-mid (Fig. 3D) blastocyst stages. From the late-mid to late blastocyst stages, the cytoplasmic F-actin signal decreased (Fig. 3B), and therefore, the correlation with nuclear YAP levels was not present. To investigate the structure of cytoplasmic F-actin in the ICM in more detail, we also performed super-resolution microscopy of late-mid blastocyst stage embryos. 3D reconstruction of the images of the ICM region revealed the presence of strong cytoplasmic F-actin signals in the SOX2-positive ICM cells (Fig. 3E, left). In these cells, strong F-actin signals visualized the cells’ morphologies. Closer examination of the 3D images inside a cell revealed that the cytoplasmic F-actin signals were fine fibrous structures oriented towards the plasma membrane and nucleus (Fig. 3E, right).

**Figure 3.**
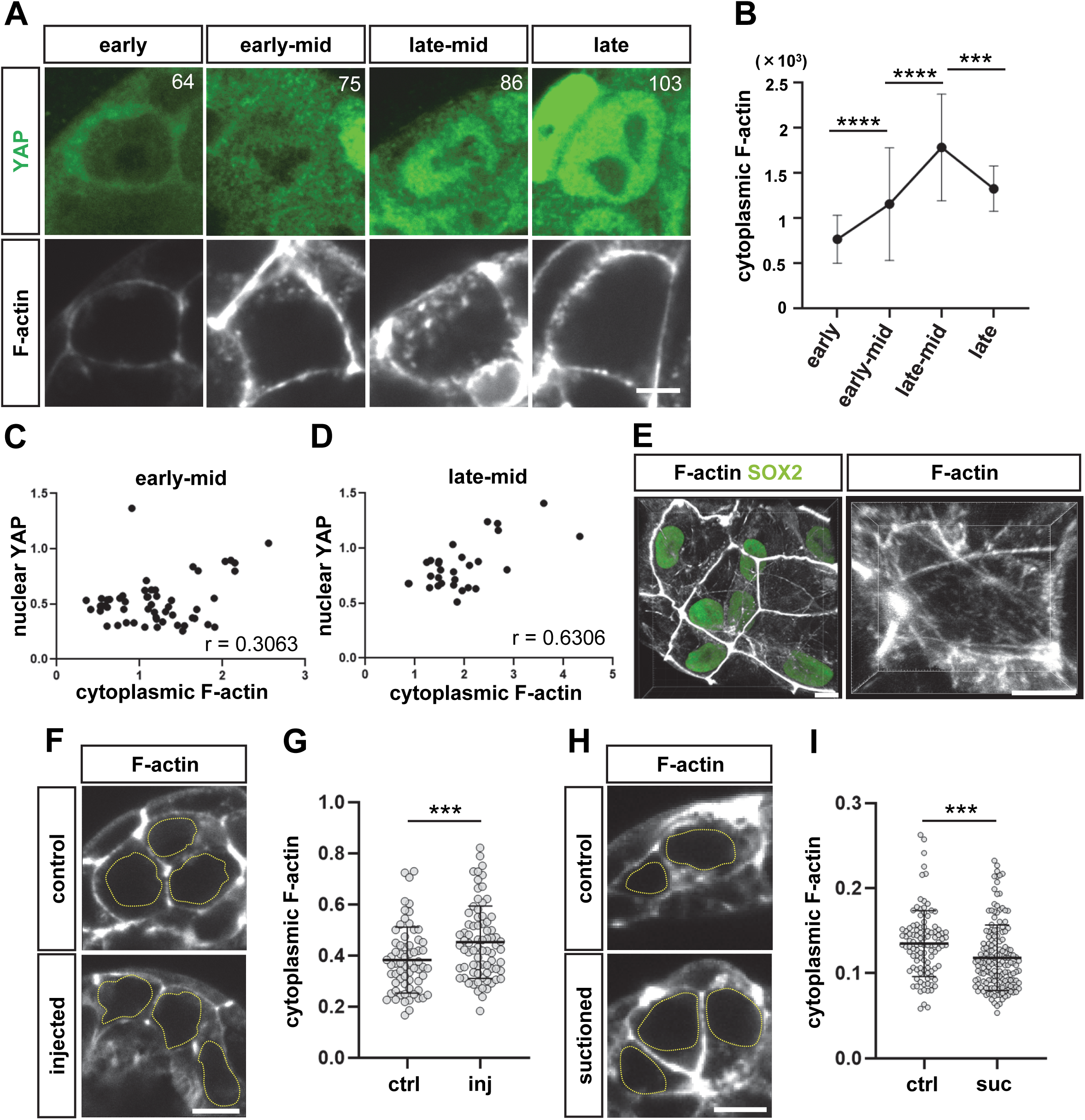
Cytoplasmic F-actin correlates with nuclear YAP in the mid blastocyst ICM. (A) Representative immunofluorescence images showing changes in YAP distribution and cytoplasmic F-actin during the blastocyst stages. The total number of cells in the embryo is indicated in the image. Dotted lines represent nuclei. Scale bar represents 5 µm. (B) A line graph showing the changes in the cytoplasmic F-actin signal during the blastocyst stages. The values shown represent the means and standard errors. *p*-values were determined by one-way ANOVA, followed by Dunn’s multiple comparison test. \*\*\**p* < 0.001, \*\*\*\**p* < 0.0001. Sample numbers analyzed for each stage were as follows: early (N = 6, n = 84), early-mid (N = 6, n = 75), late-mid (N = 5, n = 37), and late (N = 4, n = 32) blastocyst. (C, D) Dot plots showing correlations between cytoplasmic F-actin/phalloidin and nuclear YAP signals in the ICM of early-mid (C) and late-mid (D) blastocyst stage embryos. Sample numbers analyzed for each stage were as follows: early-mid (n = 56) and late-mid (n = 29) blastocyst. (E) Representative immunofluorescence super-resolution images of F-actin/phalloidin, SOX2, and nuclear YAP signals in the ICM. The left panel shows a wide area. Strong web-like F-actin signals are apical junctions of the trophectoderm cells overlying the ICM. The right panel shows an enlarged image of the inside of an ICM cell. Scale bars represent 10 μm. (F) Representative immunofluorescence images of the cytoplasmic F-actin in the ICM of blastocoel expanded embryos. Yellow dotted lines indicate nuclei. Scale bar represents 10 μm. (G) Dot plots showing increase in cytoplasmic F-actin signals in blastocoel expanded embryos. Horizontal lines represent the means and standard errors. *p*-values were determined by Student’s *t*-test. \*\*\**p* < 0.001. Sample numbers analyzed for each group were as follows: control (n = 66), injected (n = 76). (H) Representative immunofluorescence images of the cytoplasmic F-actin in the ICM of blastocoel shrunk embryos. Yellow dotted lines indicate nuclei. Scale bar represents 10 μm. (I) Dot plots showing decrease in cytoplasmic F-actin signals in blastocoel shrunk embryos. Horizontal lines represent the means and standard errors. *p*-values were determined by Student’s *t*-test. \*\*\**p* < 0.001. Sample numbers analyzed for each group were as follows: control (n = 105), suctioned (n = 162).

Correlation between cytoplasmic F-actin and nuclear YAP was also observed in blastocoel-manipulated embryos. The level of cytoplasmic F-actin signal intensity was significantly increased in blastocoel-expanded embryos (Fig. 3F, G), whereas it was significantly decreased in blastocoel-reduced embryos (Fig. 3H, I). Taken together, the amount of cytoplasmic F-actin correlates with the nuclear localization of YAP.

To examine whether cytoplasmic F-actin plays a role in YAP regulation, we experimentally manipulated F-actin. Since pharmacological disruption of F-actin caused blastocoel shrinkage and strongly reduced the viability of embryos (data not shown), we took an alternative approach. Expansion of the blastocoel alters cell shape by applying mechanical forces to the plasma membrane, and mechanical forces of the plasma membrane are transmitted to the nucleus through cytoplasmic F-actin. In cultured cells, cytoplasmic F-actin transmits the force to the nuclear lamina through interaction with a linker of nucleoskeleton and cytoskeleton (LINK) complex (Kirby and Lammerding, 2018). The nuclear lamina consists of lamin intermediate filaments and their associated proteins. The lamin family includes LAMIN A/C and LAMIN B1/2 (Dittmer and Misteli, 2011), and the amount of cytoplasmic F-actin and LAMIN A/C changes depending on the cell’s tension. The cells that receive stronger tension have more cytoplasmic F-actin and LAMIN A/C (Buxboim et al., 2014; Swift et al., 2013). We hypothesized that the cytoplasmic F-actin in ICM cells plays a similar role. Therefore, to reduce cytoplasmic F-actin, we disrupted the *Lmna* gene encoding LAMIN A/C using genome editing. One-cell stage embryos were introduced with CRISPR/Cas9 and two *Lmna* sgRNAs via electroporation (Fig. 4A). The efficiency of *Lmna* disruption was 55% (n = 15/27) as confirmed by the absence of LAMIN A/C proteins with immunostaining (Fig. 4B, C). The embryos lacking LAMIN A/C proteins were used for the analysis (Fig. 4D, E). The *Lmna* Crispants showed no obvious abnormalities in cell differentiation, blastocoel expansion, and nuclear morphology (Fig. S4A-E). *Lmna* Crispants showed a significant reduction in cytoplasmic F-actin (Fig. 4F, G) and also showed a significant reduction in nuclear YAP signal levels at the late-mid blastocyst stage (Fig. 4H, I). These results are consistent with the hypothesis that cytoplasmic F-actin is required for the nuclear localization of YAP.

**Figure 4.**
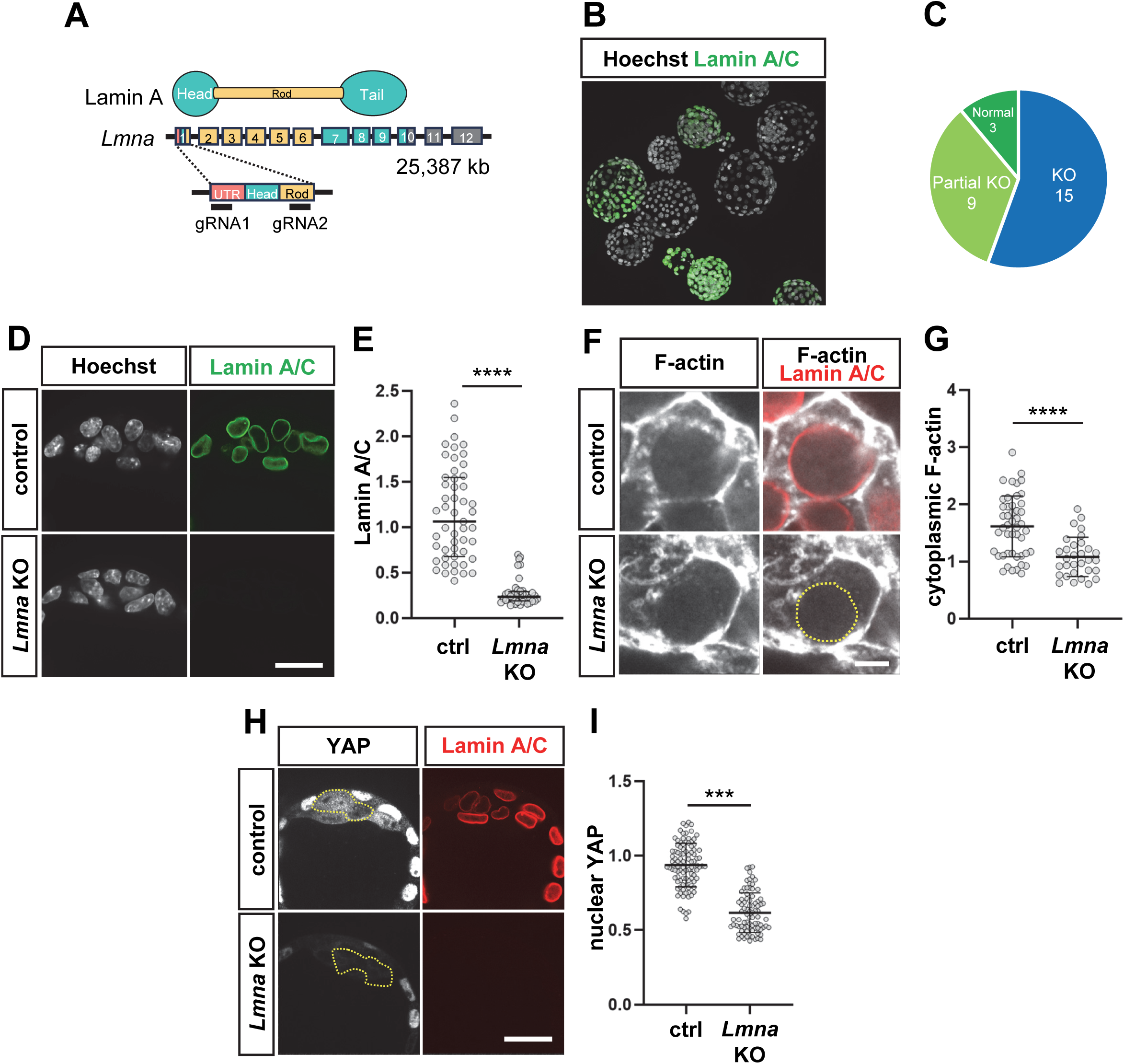
Cytoplasmic F-actin is required for nuclear YAP. (A) Schematic representation of the positions of guide RNAs for the *Lmna* gene. Top: domain structure of LAMIN A protein, middle: gene structure, bottom: enlargement of exon 1. Exons are color-coded with protein domains. (B) Representative immunofluorescence images showing reduction of LAMIN A/C expression in genome-edited embryos. Green and cyan represent LAMIN A/C and Hoechst signals, respectively. (C) A pie chart showing the knockout (KO) efficiency of the *Lmna* gene with genome editing. KO represents the embryos with no LAMIN A/C signals. Only the KO embryos were used for analyses. (D) Representative immunofluorescence images showing the absence of LAMIN A/C in *Lmna* KO embryos. Scale bar represents 25 μm. (E) Dot plots showing reduction of LAMIN A/C signals in *Lmna* KO embryos. *p*-values were determined by Student’s *t*-test. \*\*\*\**p* < 0.0001. Sample numbers analyzed for each group were as follows: control (n = 51), *Lmna* KO (n = 40). (F) Representative immunofluorescence images of the cytoplasmic F-actin in *Lmna* KO embryos. Scale bar represents 5 μm. (G) Dot plots showing decrease in cytoplasmic F-actin signals in *Lmna* KO embryos. Horizontal lines represent the means and standard errors. *p*-values were determined by Student’s *t*-test. \*\*\*\**p* < 0.0001. Sample numbers analyzed for each group were as follows: control (n = 47), *Lmna* KO (n = 31). (H) Representative immunofluorescence images of YAP in *Lmna* KO embryos. Scale bar represents 25 μm. (I) Dot plots showing decrease in nuclear YAP signals in *Lmna* KO embryos. Horizontal lines represent the means and standard errors. *p*-values were determined by Student’s *t*-test. \*\*\**p* < 0.001. Sample numbers analyzed for each group were as follows: control (n = 99), *Lmna* KO (n = 84).

### Tankyrase-dependent degradation of AMOT promotes nuclear localization of YAP

The Hippo signaling pathway plays important roles in the regulation of YAP. Activation of the Hippo pathway promotes the phosphorylation of YAP, including the serine residue 112 (S112), by LATS1/2 (Zhao et al., 2007). Therefore, we next examined the distribution of phosphorylated S112 YAP (p-YAP) to understand the changes in Hippo signaling during the blastocyst stages. The p-YAP signal tended to decrease as development progressed, with a strong reduction observed between the late-mid and late blastocyst stages (Figs. 5A, B; S1D). In this study, we used a milder fixation method to increase sensitivity, revealing the presence of p-YAP signals in the nuclei, which differs from our previous observations (Hirate et al., 2013; Nishioka et al., 2009). Consistent with previous observations, p-YAP signals were absent in the ICM of the late blastocyst stage (Hashimoto and Sasaki, 2019). These results suggest that the activity of the Hippo signaling pathway in the ICM gradually decreases during the blastocyst stages.

**Figure 5.**
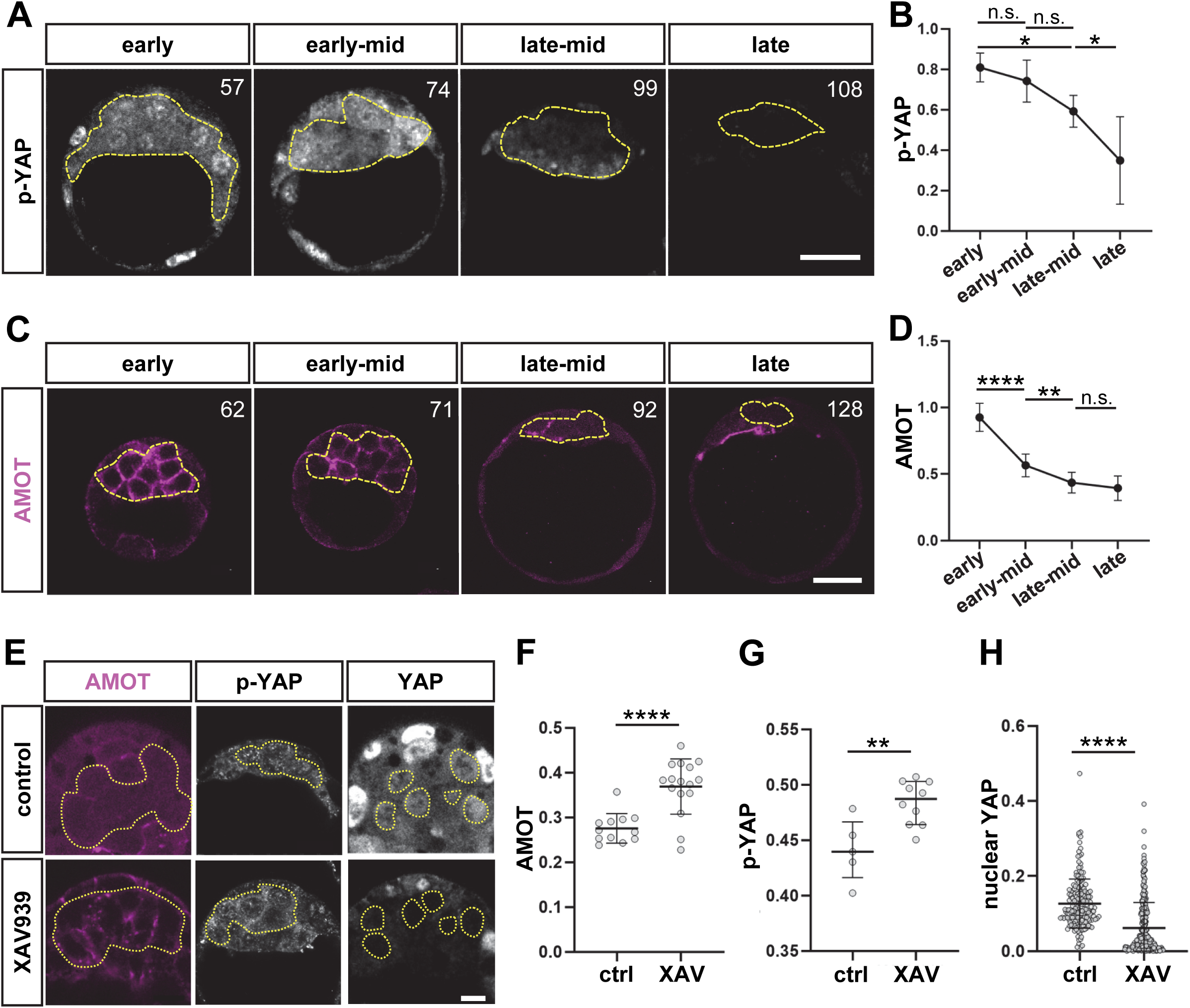
Reduction of AMOT contributes reduction of Hippo signaling in the ICM. (A) Representative immunofluorescence images of p-YAP signals during the blastocyst stage. The total number of cells in the embryo is indicated in the image. Dotted lines represent ICM/epiblast. Scale bar represents 25 μm. (B) A line graph showing the changes in the p-YAP signal during the blastocyst stages. The values shown represent the means and standard errors. *p*-values were determined by one-way ANOVA, followed by Dunn’s multiple comparison test. \**p* < 0.05, n.s.: not significant. Sample numbers analyzed for each stage were as follows: early (N = 9), early-mid (N = 5), late-mid (N = 6), and late (N = 7) blastocyst. (C) Representative immunofluorescence images of AMOT expression during the blastocyst stage. The total number of cells in the embryo is indicated in the image. Dotted lines represent ICM/epiblast. Scale bar represents 25 μm. (D) A line graph showing the changes in the AMOT signals during the blastocyst stages. The values shown represent the means and standard errors. *p*-values were determined by one-way ANOVA, followed by Dunn’s multiple comparison test. \*\**p* < 0.01, \*\*\*\**p* < 0.0001, n.s.: not significant. Sample numbers analyzed for each stage were as follows: early (N = 8), early-mid (N = 8), late-mid (N = 9), and late (N = 9) blastocyst. (E) Representative immunofluorescence images of AMOT, p-YAP and YAP signals in XAV939-treated embryos. Scale bar represents 10 μm. (F) Dot plots showing the increase in AMOT signals in XAV939-treated embryos. Horizontal lines represent the means and standard errors. *p*-values were determined by Student’s *t*-test. \*\*\*\**p* < 0.0001. Number of embryos analyzed for each group was as follows: control (N = 12), XAV939 (N = 16). (G) Dot plots showing the increase in p-YAP signals in XAV939-treated embryos. Horizontal lines represent the means and standard errors. *p*-values were determined by Student’s *t*-test. \*\**p* < 0.01. Number of embryos analyzed for each group was as follows: control (N = 5), XAV939 (N = 10). (H) Dot plots showing the decrease in nuclear YAP signals in XAV939-treated embryos. Horizontal lines represent the means and standard errors. *p*-values were determined by Student’s *t*-test. \*\*\*\**p* < 0.0001. Number of cells analyzed for each group was as follows: control (n = 186), XAV939 (n = 278).

In the ICM of early blastocyst stage embryos, Hippo signaling is activated by adherens junctions, and the association of a Hippo pathway protein, AMOT, with a junctional complex is required for activation (Hirate et al., 2013). Thus, we examined the distribution of AMOT proteins in the ICM during the blastocyst stage. A strong AMOT signal was observed at all junctions in the early blastocyst stage (Fig. 5C), as reported previously (Hirate et al., 2013; Leung and Zernicka-Goetz, 2013). Then, the AMOT signal was strongly reduced in the early-mid blastocyst stage and was observed only at some cell junctions in the late-mid blastocyst stage (Fig. 5C, D). In late blastocysts, no AMOT signal was observed in the SOX2-positive epiblast cells, similar to our previous observation (Figs. 5C, D; S1E) (Hashimoto and Sasaki, 2019). Thus, the AMOT signal in the blastocyst stage gradually decreases during differentiation.

We next asked whether the reduction of AMOT is responsible for the increase in nuclear YAP levels. In cultured cells, AMOT family proteins are subject to ubiquitin-dependent degradation through poly-ADP ribosylation by tankyrases (TNKS1 and TNKS2), followed by ubiquitination with RNF146 E3 ligase. Treatment with a tankyrase inhibitor, XAV939, stabilizes AMOT (Wang et al., 2015). To examine whether a tankyrase-dependent degradation mechanism is involved in the reduction of AMOT in the ICM, we treated early-mid blastocyst stage embryos with XAV939 for 12 hours until the late-mid blastocyst stage. XAV939-treated embryos had a comparable number of cells to control embryos, indicating that the treatment did not have significant effects on development (Fig. S5A). In the ICM of XAV939-treated embryos, AMOT signal was significantly increased (Fig. 5E, F), suggesting that a tankyrase-dependent degradation is a key mechanism of AMOT degradation in the ICM of mid-blastocyst stage embryos. In these XAV939-treated AMOT-upregulated embryos, p-YAP signal was significantly increased, and nuclear YAP signal was significantly reduced (Fig. 5E, G, H). Taken together, these results suggest that the reduction of AMOT proteins by a tankyrase-dependent degradation mechanism contributes to the nuclear localization of YAP by inactivating Hippo signaling at the mid blastocyst stage.

### SOX2 is required for nuclear localization of YAP

Hippo signaling and YAP localization in the ICM gradually change during the blastocyst stage. This prompted us to hypothesize that the specification and/or differentiation of the ICM alters Hippo/YAP regulation. To test this hypothesis, we focused on the transcription factor SOX2 because it is the first transcription factor specifically expressed in the inner cells of the morula and the ICM of early blastocysts before YAP nuclear localization (Guo et al., 2010; Wicklow et al., 2014). The expression of SOX2 rises rapidly from the early-mid blastocyst stage (Fig. 6A, B). To examine the role of SOX2, we generated *Sox2* Crispant embryos by genome editing of zygotes. The efficiency of *Sox2* disruption, confirmed by the absence of SOX2 signal, is 100% (N = 50/50 embryos) (Fig. 6C, D). At the mid blastocyst stage, *Sox2* Crispant embryos showed significantly lower nuclear YAP signals (Fig. 6E, F), indicating that SOX2 is required for nuclear localization of YAP. Furthermore, AMOT and p-YAP signals were significantly elevated in *Sox2* Crispant embryos (Fig. 6G-J). These results suggest that SOX2 promotes nuclear localization of YAP via downregulation of AMOT and Hippo signaling. Downregulation of AMOT likely involves a tankyrase-dependent degradation mechanism, as shown above.

**Figure 6.**
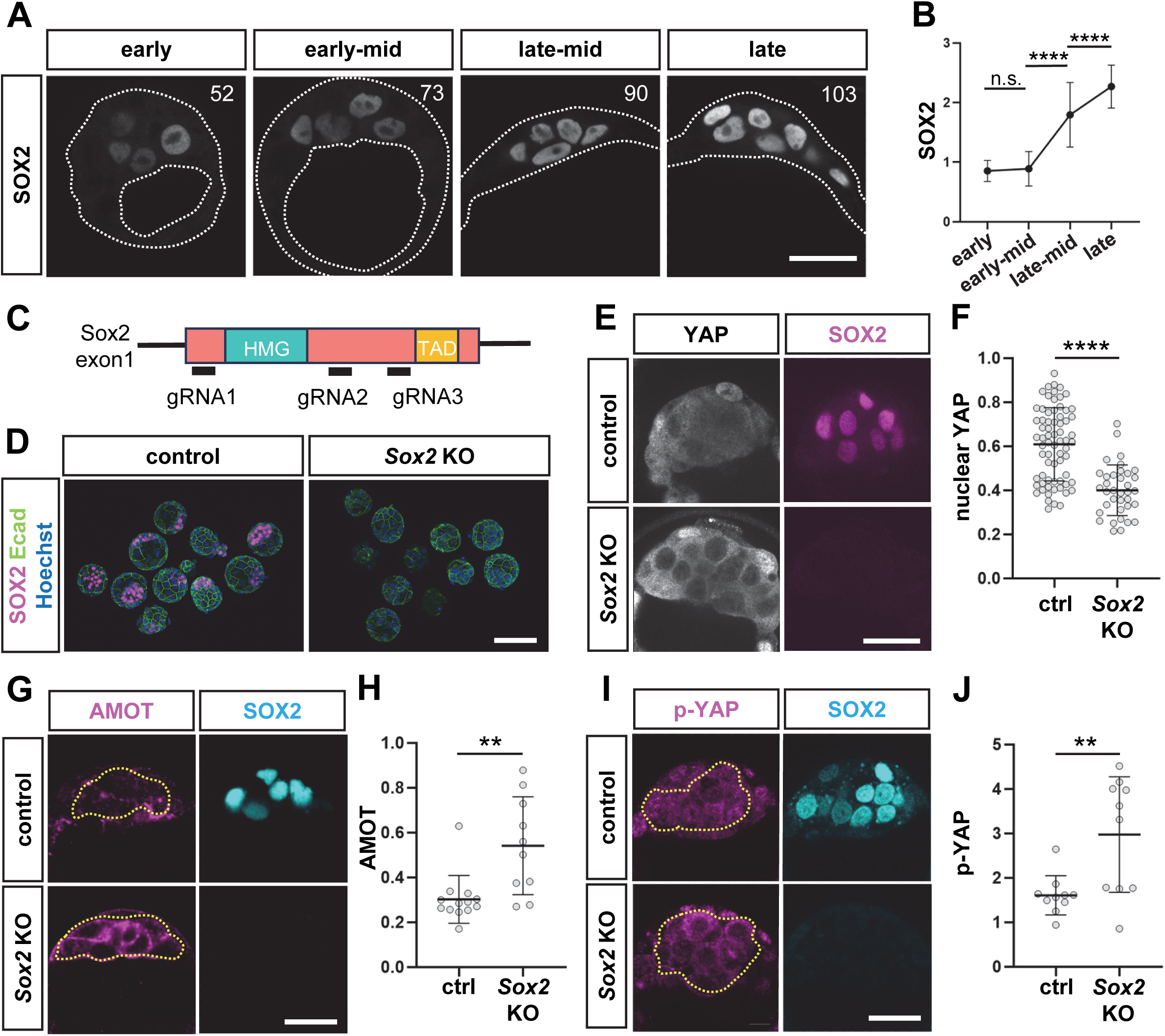
SOX2 is required for nuclear translocation of YAP in the ICM. (A) Representative immunofluorescence images of SOX2 expression during the blastocyst stage. The total number of cells in the embryo is indicated in the image. Dashed lines represent embryo and blastocoel. Scale bar represents 25 μm. (B) A line graph showing the changes in the SOX2 signals during the blastocyst stages. The values shown represent the means and standard errors. *p*-values were determined by one-way ANOVA, followed by Dunn’s multiple comparison test. \*\*\*\**p* < 0.0001, n.s.: not significant. Number of cells analyzed for each stage was as follows: early (n = 109), early-mid (n = 70), late-mid (n = 67), and late (n = 118) blastocyst. (C) Schematic representation of the positions of guide RNAs for the *Sox2* gene. Genomic organization is shown. The relationship to the protein domains is shown. (D) Representative immunofluorescence images showing absence of SOX2 expression in *Sox2* Crispant embryos. Scale bar represents 100 μm. (E) Representative immunofluorescence images of YAP and SOX2 expression in *Sox2* KO embryos. Scale bar represents 25 μm. (F) Dot plots showing the decrease in nuclear YAP signals in *Sox2* KO embryos. Horizontal lines represent the means and standard errors. *p*-values were determined by Student’s *t*-test. \*\*\*\**p* < 0.0001. Number of cells analyzed for each group was as follows: control (n = 69), *Sox2* KO (n = 36). (G) Representative immunofluorescence images of AMOT and SOX2 expression in *Sox2* KO embryos. Dotted lines delineate ICM. Scale bar represents 25 μm. (H) Dot plots showing the increase in AMOT signals in *Sox2* KO embryos. Horizontal lines represent the means and standard errors. *p*-values were determined by Student’s *t-*test. \*\**p* < 0.01. Number of embryos analyzed for each group was as follows: control (N = 10), *Sox2* KO (N = 10). (I) Representative immunofluorescence images of p-YAP and SOX2 expression in *Sox2* KO embryos. Dotted lines represent ICM. Scale bar represents 25 μm. (J) Dot plots showing the increase in p-YAP signals in *Sox2* KO embryos. Horizontal lines represent the means and standard errors. *p*-values were determined by Student’s *t*-test. \*\**p* < 0.01. Number of embryos analyzed for each group was as follows: control (N = 13), *Sox2* KO (N = 10).

### Blastocoel expansion influences YAP localization independent of AMOT-mediated Hippo signaling

We have shown the operation of two distinct mechanisms promoting nuclear localization of YAP in the ICM: a mechano-physical control by the expansion of the blastocoel and a molecular control of Hippo signaling triggered by junctional AMOT. To test the relationship between these two mechanisms, we asked whether YAP phosphorylation and AMOT expression were affected by manipulations of blastocoel size. In blastocoel-expanded embryos, the p-YAP signal was significantly reduced, indicating a reduction of Hippo signaling (Fig. 7A, B). In blastocoel-reduced embryos, the p-YAP signal was significantly increased, indicating activation of Hippo signaling (Fig. 7C, D). However, manipulations of blastocoel size did not significantly alter the AMOT signals (Fig. 7E-H), suggesting that mechano-physical status controls YAP phosphorylation independent of AMOT. Taken together, these results suggest that both mechano-physical and molecular mechanisms regulate YAP localization through phosphorylation, but these mechanisms act in parallel upstream of YAP phosphorylation.

**Figure 7.**
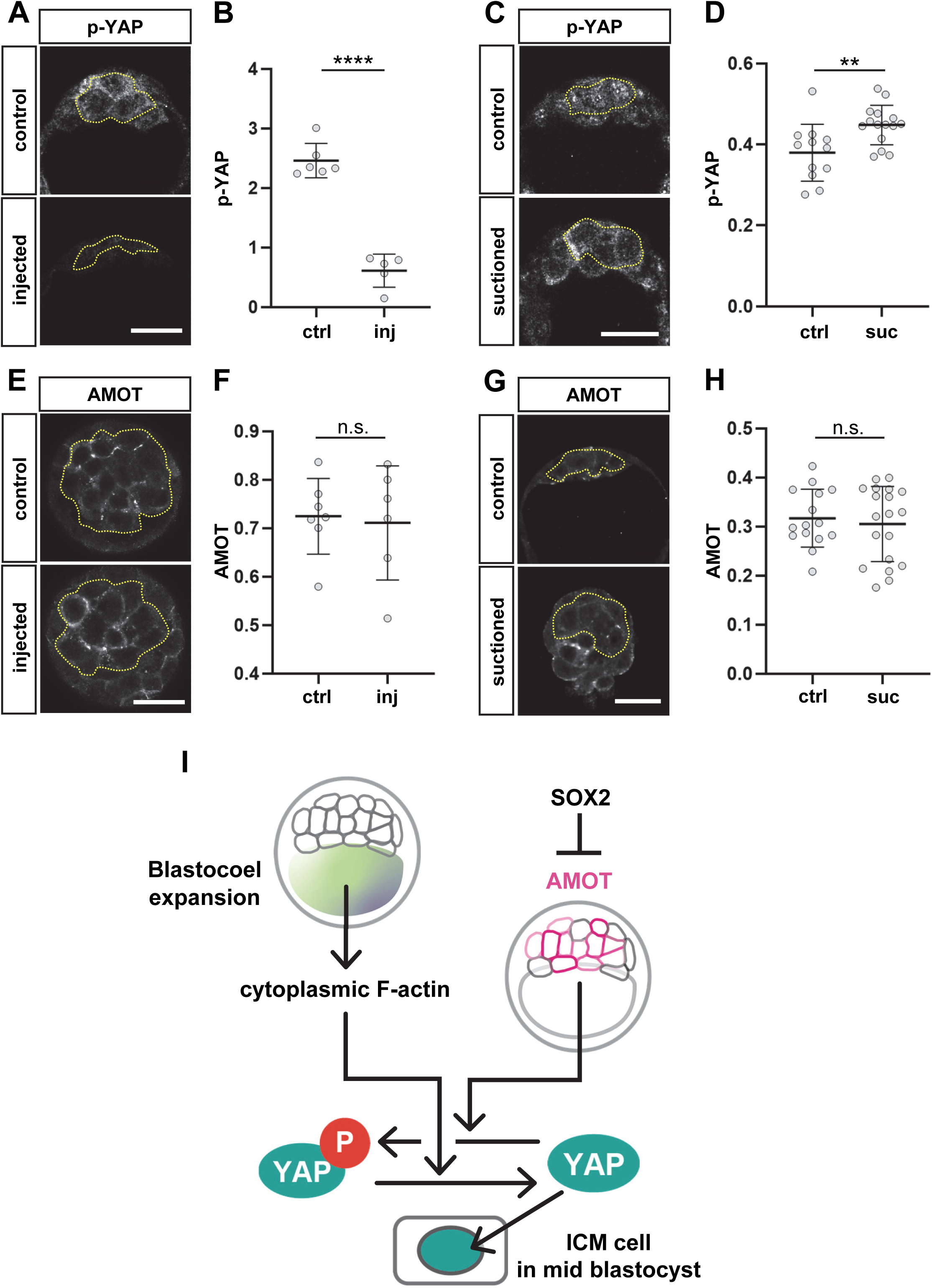
Blastocoel size manipulation alters Hippo signaling without affecting AMOT. (A) Representative immunofluorescence images of p-YAP signals in blastocoel expanded embryos. Dotted lines represent ICM. Scale bar represents 25 μm. (B) Dot plots showing the decrease in p-YAP signals in blastocoel expanded embryos. Horizontal lines represent the means and standard errors. *p*-values were determined by Student’s *t*-test. \*\*\*\**p* < 0.0001. Number of embryos analyzed for each group was as follows: control (N = 6), injected (N = 5). (C) Representative immunofluorescence images of p-YAP signals in blastocoel sucked embryos. Dotted lines represent ICM. Scale bar represents 25 μm. (D) Dot plots showing the increase in p-YAP signals in blastocoel sucked embryos. Horizontal lines represent the means and standard errors. *p*-values were determined by Student’s *t*-test. \*\**p* < 0.01. Number of embryos analyzed for each group was as follows: control (N = 12), suctioned (N = 15). (E) Representative immunofluorescence images of AMOT signals in blastocoel expanded embryos. Dotted lines represent ICM. Scale bar represents 25 μm. (F) Dot plots showing no effect on AMOT expression in blastocoel expanded embryos. Horizontal lines represent the means and standard errors. *p*-values were determined by Student’s *t*-test. n.s.; not significant. Number of embryos analyzed for each group was as follows: control (N = 7), injected (N = 6). (G) Representative immunofluorescence images of AMOT signals in blastocoel sucked embryos. Dotted lines represent ICM. Scale bar represents 25 μm. (H) Dot plots showing no effect on AMOT expression in blastocoel sucked embryos. Horizontal lines represent the means and standard errors. *p*-values were determined by Student’s *t*-test. n.s.; not significant. Number of embryos analyzed for each group was as follows: control (N = 15), suctioned (N = 19). (I) Model of YAP regulation in the ICM/epiblast during epiblast formation.

## Discussion

In this study, we examined the mechanisms promoting nuclear localization of YAP in the ICM and epiblast during blastocyst development. Based on our results and previous findings, we propose a model of YAP regulation during epiblast formation (Fig. 7I).

At the early blastocyst stage, AMOT is present at adherens junctions, activating the Hippo signaling pathway (Hirate et al., 2013). Active Hippo signaling promotes the phosphorylation of YAP, resulting in its cytoplasmic localization. During the mid-blastocyst stage, the expansion of the blastocoel exerts force on the trophectoderm, increasing its cortical tension (Chan et al., 2019). A similar force is likely applied to the ICM. ICM cells experiencing this physical force increase their cytoplasmic F-actin levels. This increase in cytoplasmic F-actin reduces YAP phosphorylation, promoting its nuclear localization. Cytoskeleton-mediated regulation of YAP involves both LATS-dependent (Meng et al., 2018; Rauskolb et al., 2014; Wada et al., 2011; Zhao et al., 2012) and independent mechanisms (Aragona et al., 2013; Chang et al., 2018; Dupont et al., 2011; Elosegui-Artola et al., 2017). Our results suggest that mechanical cues regulate YAP through LATS-dependent mechanisms in forming epiblast cells.

In parallel, differentiation of ICM/epiblast cells also contributes to YAP nuclear localization. In the ICM of early blastocysts, junctional AMOT proteins activate Hippo signaling and suppress nuclear localization of YAP. During the blastocyst stage, junctional AMOT gradually reduces, diminishing Hippo pathway activation and promoting YAP nuclear localization. The reduction of AMOT at the mid-blastocyst stage is caused by post-transcriptional degradation through tankyrase- and ubiquitin-dependent mechanisms (Wang et al., 2015), as inhibition of tankyrase activity significantly increased AMOT. SOX2 regulates the reduction of AMOT proteins. SOX2 is the first transcription factor specifically expressed in the inner cells of the morula and the ICM of early blastocysts (Guo et al., 2010; Wicklow et al., 2014), and it is essential for ICM specification and development, or the acquisition of a pre-pluripotency state (Li et al., 2023; Wicklow et al., 2014). In the absence of SOX2, the ICM maintains strong expression of AMOT and cytoplasmic localization of YAP with active Hippo signaling, similar to early ICM conditions.

Although both mechano-physical (blastocoel–F-actin) and cell differentiation (SOX2–AMOT) mechanisms regulate YAP localization through its phosphorylation, they function in parallel, as the former mechanism did not alter the amount of junctional AMOT proteins. Thus, the mechano-physical mechanism directly suppresses LATS1/2 downstream of F-actin to inhibit YAP phosphorylation, while the cell differentiation mechanism reduces Hippo pathway activation at adherens junctions by decreasing AMOT proteins. The detailed mechanisms of these pathways, such as the regulation of LATS1/2 by F-actin and tankyrase activities by SOX2, remain to be elucidated in future studies.

At the mid-blastocyst stage, AMOT is primarily regulated by post-transcriptional degradation. In the epiblast of late blastocysts, however, *Amot* RNA is not expressed (Guo et al., 2010; Nakamura et al., 2015; Stirparo et al., 2021), suggesting that AMOT is also transcriptionally regulated at the late stage. Therefore, in the transitional state of the mid-blastocyst stage, post-transcriptional regulation of AMOT rapidly modulates YAP to control epiblast differentiation. Upon epiblast differentiation, Hippo signaling is inactivated at the transcriptional level to maintain the epiblast state. Mechanical mechanisms predominantly contribute to the early to mid-blastocyst stages, as the blastocoel undergoes cyclic oscillation at the late blastocyst stage (Chan et al., 2019).

Our previous study showed that variation in nuclear YAP levels among ICM cells at the mid-blastocyst stage triggers cell competition (Hashimoto and Sasaki, 2019). One question that arises is what causes the variation in nuclear YAP levels. Our study revealed that both mechano-physical and cell differentiation mechanisms regulate YAP. A strong correlation between cytoplasmic F-actin and nuclear YAP suggests that variations in the mechanical forces that cells receive contribute to the variation in nuclear YAP levels. Despite the strong correlation between cytoplasmic F-actin and nuclear YAP, the correlation between nuclear shape and nuclear YAP is low. Thus, variations in the mechanical forces experienced by cells are probably not strong enough to significantly alter nuclear shape but are sufficient to affect F-actin levels. Considering that the timing of epiblast fate specification varies among ICM cells (Saiz et al., 2016), variation in cell differentiation also contributes to the differences in nuclear YAP levels.

## Conclusion

In conclusion, we have shown that mechano-physical and cell differentiation mechanisms cooperatively regulate the nuclear localization of YAP. The contribution of these mechanisms may vary depending on the developmental stage. The operation of these two mechanisms likely enhances the robustness of epiblast differentiation.

## Acknowledgments

We thank Dr. Masakazu Hashimoto for his advice on embryo manipulations and discussions, Dr. Syuichi Onami and Dr. Yusuke Azuma for their advice on quantitative image analysis at the outset of this project, Dr. Feng Zhang and Addgene for providing the pX330 plasmid. We also thank Daisuke Masui, Rui Shibata, Kanta Arai, Megumi Yokoshima, and the Animal Facility of the Frontier of Biosciences at the University of Osaka for their dedicated efforts in maintaining the mouse colonies essential for this study.

## Author contributions

Conceptualization: H.S.; Methodology: H.S, H.M.; Validation: H.S.; Formal analysis: H.M.; Investigation: H.M.; Data curation: H.M., H.S.; Writing - original draft: H.M., H.S.; Writing - review & editing: H.S.; Visualization: H.M.; Supervision: H.S.; Project administration: H.S.; Funding acquisition: H.M., H.S.

## Declaration of Generative AI and AI-assisted technologies in the writing process

During the preparation of this work, the authors used ChatGPT in order to check grammar and improve readability of the text. After using this tool, the authors reviewed and edited the content as needed and take full responsibility for the content of the publication.

## Legends for Supplementary Figures

**Figure S1. Relationship between total cell number and various parameters**

(A) A dot plot showing the relationship between total cell number and blastocoel volume. Each dot represents one embryo.

(B) A dot plot showing the relationship between total cell number and sphericity of the nucleus in the ICM/epiblast.

(C) A dot plot showing the relationship between total cell number and cytoplasmic F-actin signals in the ICM/epiblast.

(D) A dot plot showing the correlation between total cell number and p-YAP signals in the ICM/epiblast.

(E) A dot plot showing the correlation between total cell number and AMOT signals in the ICM/epiblast.

**Figure S2. Blastocoel expansion showed increased YAP N/C ratio.**

(A) Representative images of the segmentations of nuclei and cytoplasm in the ICM used for quantification of YAP N/C ratio. Red and white represent nuclei and cytoplasm. Only a small portion of cytoplasm was segmented for each cell in this analysis. Scale bar represents 20 μm.

(B) Dot plots showing the increase in YAP N/C ratio with blastocoel expansion. Horizontal lines represent the means and standard errors. *p*-values were determined by Student’s *t*-test. **p < 0.01. Sample number analyzed for each group was as follows: control (n = 88), blastocoel expansion (n = 93).

**Figure S3. Relationship between nuclear morphology and nuclear YAP**

(A) Representative images of surface models of the ICM/epiblast nuclei showing sphericity changes during the blastocyst stages. Scale bar represents 15 μm.

(B) A line graph showing the changes in sphericity during the blastocyst stages. The values shown represent the means and standard errors. *p*-values were determined by one-way ANOVA, followed by Dunn’s multiple comparison test. \**p* < 0.05, n.s.: not significant. Sample number analyzed for each stage was as follows: early (N = 10), early-mid (N = 15), late-mid (N = 16), and late (N = 16) blastocyst.

(C, D) Dot plots showing the relationship between nuclear sphericity and nuclear YAP signals in the ICM of early-mid (C) and late-mid (D) blastocyst stage embryos. Sample number analyzed for each stage was as follows: early-mid (n = 263) and late-mid (n = 216) blastocyst.

**Figure S4. *Lmna* knockout embryos showed no obvious abnormalities**

(A) Representative immunofluorescence images of Hoechst signals in control and *Lmna* KO embryos.

(B) Representative immunofluorescence images of Hoechst and SOX2 signals in control and *Lmna* KO embryos. Scale bar represents 25 μm.

(C) Representative immunofluorescence images of Hoechst and SOX17 signals in control and *Lmna* KO embryos. Scale bar represents 25 μm.

(D) Representative images of surface models of the ICM/epiblast nuclei in control and *Lmna* KO embryos. Scale bar represents 15 μm.

(E) Dot plots showing similar sphericities between control and *Lmna* KO embryos. Horizontal lines represent the means and standard errors. *p*-values were determined by Student’s *t*-test. n.s.; not significant. Sample number analyzed for each group was as follows: control (n = 99), *Lmna* KO (n = 100).

**Figure S5. XAV939 treatment does not affect cell numbers.**

(A) Dot plots showing similar cell numbers in XAV939 treated embryos. Horizontal lines represent the means and standard errors. *p*-values were determined by Student’s *t*-test. n.s.; not significant. Number of embryos analyzed for each group was as follows: control (N = 12), XAV939 (N = 16).

**Figure S6. Segmentation methods used for quantification of signals**

(A) A representative image of a surface model of the ICM/epiblast nuclei used for quantification of nuclear signals.

(B) Representative images of segmentation of the cytoplasmic region in one optical plane (left) and a surface model of a cytoplasmic region (right) used for quantification of cytoplasmic signals. Scale bar represents 10 μm.

(C) A representative image of segmentation of the ICM/epiblast region in one optical plane used for quantification of p-YAP signals. Scale bar represents 10 μm.

(D) Representative images of segmentation of the ICM/epiblast region in one optical plane (left) and a surface model of an ICM/epiblast region (right) used for quantification of AMOT signals. Scale bar represents 10 μm.

## References

1. Anani, S., Bhat, S., Honma-Yamanaka, N., Krawchuk, D., Yamanaka, Y., 2014. Initiation of Hippo signaling is linked to polarity rather than to cell position in the pre-implantation mouse embryo. Development (Cambridge, England) 141, 2813–2824.

2. Aragona, M., Panciera, T., Manfrin, A., Giulitti, S., Michielin, F., Elvassore, N., Dupont, S., Piccolo, S., 2013. A mechanical checkpoint controls multicellular growth through YAP/TAZ regulation by actin-processing factors. Cell 154, 1047–1059.

3. Buxboim, A., Swift, J., Irianto, J., Spinler, K.R., Dingal, P.C.D.P., Athirasala, A., Kao, Y.-R.C., Cho, S., Harada, T., Shin, J.-W., Discher, D.E., 2014. Matrix elasticity regulates lamin-A,C phosphorylation and turnover with feedback to actomyosin. Current biology : CB 24, 1909–1917.

4. Chan, C.J., Costanzo, M., Ruiz-Herrero, T., Monke, G., Petrie, R.J., Bergert, M., Diz-Munoz, A., Mahadevan, L., Hiiragi, T., 2019. Hydraulic control of mammalian embryo size and cell fate. Nature 571, 112–116.

5. Chan, S.W., Lim, C.J., Loo, L.S., Chong, Y.F., Huang, C., Hong, W., 2009. TEADs mediate nuclear retention of TAZ to promote oncogenic transformation. The Journal of biological chemistry 284, 14347–14358.

6. Chang, L., Azzolin, L., Di Biagio, D., Zanconato, F., Battilana, G., Lucon Xiccato, R., Aragona, M., Giulitti, S., Panciera, T., Gandin, A., Sigismondo, G., Krijgsveld, J., Fassan, M., Brusatin, G., Cordenonsi, M., Piccolo, S., 2018. The SWI/SNF complex is a mechanoregulated inhibitor of YAP and TAZ. Nature 563, 265–269.

7. Cockburn, K., Rossant, J., 2010. Making the blastocyst: lessons from the mouse. The Journal of clinical investigation 120, 995–1003.

8. Cong, L., Ran, F.A., Cox, D., Lin, S., Barretto, R., Habib, N., Hsu, P.D., Wu, X., Jiang, W., Marraffini, L.A., Zhang, F., 2013. Multiplex genome engineering using CRISPR/Cas systems. Science (New York, N.Y 339, 819–823.

9. Dittmer, T.A., Misteli, T., 2011. The lamin protein family. Genome Biol 12, 222.

10. Driscoll, T.P., Cosgrove, B.D., Heo, S.-J., Shurden, Z.E., Mauck, R.L., 2015. Cytoskeletal to Nuclear Strain Transfer Regulates YAP Signaling in Mesenchymal Stem Cells. Biophysical journal 108, 2783–2793.

11. Dupont, S., Morsut, L., Aragona, M., Enzo, E., Giulitti, S., Cordenonsi, M., Zanconato, F., Le Digabel, J., Forcato, M., Bicciato, S., Elvassore, N., Piccolo, S., 2011. Role of YAP/TAZ in mechanotransduction. Nature 474, 179–183.

12. Elosegui-Artola, A., Andreu, I., Beedle, A.E.M., Lezamiz, A., Uroz, M., Kosmalska, A.J., Oria, R., Kechagia, J.Z., Rico-Lastres, P., Le Roux, A.L., Shanahan, C.M., Trepat, X., Navajas, D., Garcia-Manyes, S., Roca-Cusachs, P., 2017. Force Triggers YAP Nuclear Entry by Regulating Transport across Nuclear Pores. Cell 171, 1397–1410 e1314.

13. Frum, T., Watts, J.L., Ralston, A., 2019. TEAD4, YAP1 and WWTR1 prevent the premature onset of pluripotency prior to the 16-cell stage. Development (Cambridge, England) 146.

14. Guo, G., Huss, M., Tong, G.Q., Wang, C., Li Sun, L., Clarke, N.D., Robson, P., 2010. Resolution of cell fate decisions revealed by single-cell gene expression analysis from zygote to blastocyst. Developmental cell 18, 675–685.

15. Hashimoto, M., Sasaki, H., 2019. Epiblast Formation by TEAD-YAP-Dependent Expression of Pluripotency Factors and Competitive Elimination of Unspecified Cells. Developmental cell 50, 139–154 e135.

16. Hashimoto, M., Yamashita, Y., Takemoto, T., 2016. Electroporation of Cas9 protein/sgRNA into early pronuclear zygotes generates non-mosaic mutants in the mouse. Developmental biology 418, 1–9.

17. Hirate, Y., Hirahara, S., Inoue, K.I., Suzuki, A., Alarcon, V.B., Akimoto, K., Hirai, T., Hara, T., Adachi, M., Chida, K., Ohno, S., Marikawa, Y., Nakao, K., Shimono, A., Sasaki, H., 2013. Polarity-Dependent Distribution of Angiomotin Localizes Hippo Signaling in Preimplantation Embryos. Curr Biol 23, 1181–1194.

18. Kirby, T.J., Lammerding, J., 2018. Emerging views of the nucleus as a cellular mechanosensor. Nature cell biology 20, 373–381.

19. Labun, K., Montague, T.G., Krause, M., Torres Cleuren, Y.N., Tjeldnes, H., Valen, E., 2019. CHOPCHOP v3: expanding the CRISPR web toolbox beyond genome editing. Nucleic Acids Res 47, W171–W174.

20. Lei, Q.Y., Zhang, H., Zhao, B., Zha, Z.Y., Bai, F., Pei, X.H., Zhao, S., Xiong, Y., Guan, K.L., 2008. TAZ promotes cell proliferation and epithelial-mesenchymal transition and is inhibited by the hippo pathway. Molecular and cellular biology 28, 2426–2436.

21. Leung, C.Y., Zernicka-Goetz, M., 2013. Angiomotin prevents pluripotent lineage differentiation in mouse embryos via Hippo pathway-dependent and -independent mechanisms. Nature communications 4, 2251.

22. Li, L., Lai, F., Hu, X., Liu, B., Lu, X., Lin, Z., Liu, L., Xiang, Y., Frum, T., Halbisen, M.A., Chen, F., Fan, Q., Ralston, A., Xie, W., 2023. Multifaceted SOX2-chromatin interaction underpins pluripotency progression in early embryos. Science (New York, N.Y 382, eadi5516.

23. Ma, S., Meng, Z., Chen, R., Guan, K.-L., 2019. The Hippo Pathway: Biology and Pathophysiology. Annu Rev Biochem 88, 577–604.

24. Maitre, J.L., Turlier, H., Illukkumbura, R., Eismann, B., Niwayama, R., Nedelec, F., Hiiragi, T., 2016. Asymmetric division of contractile domains couples cell positioning and fate specification. Nature 536, 344–348.

25. Meng, Z., Qiu, Y., Lin, K.C., Kumar, A., Placone, J.K., Fang, C., Wang, K.-C., Lu, S., Pan, M., Hong, A.W., Moroishi, T., Luo, M., Plouffe, S.W., Diao, Y., Ye, Z., Park, H.W., Wang, X., Yu, F.-X., Chien, S., Wang, C.-Y., Ren, B., Engler, A.J., Guan, K.-L., 2018. RAP2 mediates mechanoresponses of the Hippo pathway. Nature 560, 655–660.

26. Nakamura, T., Yabuta, Y., Okamoto, I., Aramaki, S., Yokobayashi, S., Kurimoto, K., Sekiguchi, K., Nakagawa, M., Yamamoto, T., Saitou, M., 2015. SC3-seq: a method for highly parallel and quantitative measurement of single-cell gene expression. Nucleic Acids Res 43, e60.

27. Nishioka, N., Inoue, K., Adachi, K., Kiyonari, H., Ota, M., Ralston, A., Yabuta, N., Hirahara, S., Stephenson, R.O., Ogonuki, N., Makita, R., Kurihara, H., Morin-Kensicki, E.M., Nojima, H., Rossant, J., Nakao, K., Niwa, H., Sasaki, H., 2009. The Hippo signaling pathway components Lats and Yap pattern Tead4 activity to distinguish mouse trophectoderm from inner cell mass. Developmental cell 16, 398–410.

28. Ota, M., Sasaki, H., 2008. Mammalian Tead proteins regulate cell proliferation and contact inhibition as a transcriptional mediator of Hippo signaling. Development (Cambridge, England) 135, 4059–4069.

29. Ralston, A., Cox, B.J., Nishioka, N., Sasaki, H., Chea, E., Rugg-Gunn, P., Guo, G., Robson, P., Draper, J.S., Rossant, J., 2010. Gata3 regulates trophoblast development downstream of Tead4 and in parallel to Cdx2. Development (Cambridge, England) 137, 395–403.

30. Rauskolb, C., Sun, S., Sun, G., Pan, Y., Irvine, K.D., 2014. Cytoskeletal tension inhibits Hippo signaling through an Ajuba-Warts complex. Cell 158, 143–156.

31. Rossant, J., Tam, P.P., 2009. Blastocyst lineage formation, early embryonic asymmetries and axis patterning in the mouse. Development (Cambridge, England) 136, 701–713.

32. Saiz, N., Williams, K.M., Seshan, V.E., Hadjantonakis, A.K., 2016. Asynchronous fate decisions by single cells collectively ensure consistent lineage composition in the mouse blastocyst. Nature communications 7, 13463.

33. Skory, R.M., Moverley, A.A., Ardestani, G., Alvarez, Y., Domingo-Muelas, A., Pomp, O., Hernandez, B., Tetlak, P., Bissiere, S., Stern, C.D., Sakkas, D., Plachta, N., 2023. The nuclear lamina couples mechanical forces to cell fate in the preimplantation embryo via actin organization. Nature communications 14, 3101.

34. Stirparo, G.G., Kurowski, A., Yanagida, A., Bates, L.E., Strawbridge, S.E., Hladkou, S., Stuart, H.T., Boroviak, T.E., Silva, J.C.R., Nichols, J., 2021. OCT4 induces embryonic pluripotency via STAT3 signaling and metabolic mechanisms. Proceedings of the National Academy of Sciences of the United States of America 118.

35. Swift, J., Ivanovska, I.L., Buxboim, A., Harada, T., Dingal, P.C.D.P., Pinter, J., Pajerowski, J.D., Spinler, K.R., Shin, J.-W., Tewari, M., Rehfeldt, F., Speicher, D.W., Discher, D.E., 2013. Nuclear lamin-A scales with tissue stiffness and enhances matrix-directed differentiation. Science (New York, N Y) 341, 1240104.

36. Wada, K., Itoga, K., Okano, T., Yonemura, S., Sasaki, H., 2011. Hippo pathway regulation by cell morphology and stress fibers. Development (Cambridge, England) 138, 3907–3914.

37. Wadell, H., 1933. Sedimentation and Sedimentology. Science (New York, N.Y 77, 536–537.

38. Wang, W., Li, N., Li, X., Tran, M.K., Han, X., Chen, J., 2015. Tankyrase Inhibitors Target YAP by Stabilizing Angiomotin Family Proteins. Cell reports 13, 524–532.

39. Wicklow, E., Blij, S., Frum, T., Hirate, Y., Lang, R.A., Sasaki, H., Ralston, A., 2014. HIPPO pathway members restrict SOX2 to the inner cell mass where it promotes ICM fates in the mouse blastocyst. PLoS Genet 10, e1004618.

40. Zhao, B., Li, L., Wang, L., Wang, C.Y., Yu, J., Guan, K.L., 2012. Cell detachment activates the Hippo pathway via cytoskeleton reorganization to induce anoikis. Genes & development 26, 54–68.

41. Zhao, B., Wei, X., Li, W., Udan, R.S., Yang, Q., Kim, J., Xie, J., Ikenoue, T., Yu, J., Li, L., Zheng, P., Ye, K., Chinnaiyan, A., Halder, G., Lai, Z.C., Guan, K.L., 2007. Inactivation of YAP oncoprotein by the Hippo pathway is involved in cell contact inhibition and tissue growth control. Genes & development 21, 2747–2761.

42. Zhao, B., Ye, X., Yu, J., Li, L., Li, W., Li, S., Yu, J., Lin, J.D., Wang, C.Y., Chinnaiyan, A.M., Lai, Z.C., Guan, K.L., 2008. TEAD mediates YAP-dependent gene induction and growth control. Genes & development 22, 1962–1972.

43. Zheng, Y., Pan, D., 2019. The Hippo Signaling Pathway in Development and Disease. Developmental cell 50, 264–282.

