## Supplementary figures and images for "Blastocoel expansion and AMOT degradation cooperatively promote YAP nuclear localization during epiblast formation"

### Fig. S1-S6

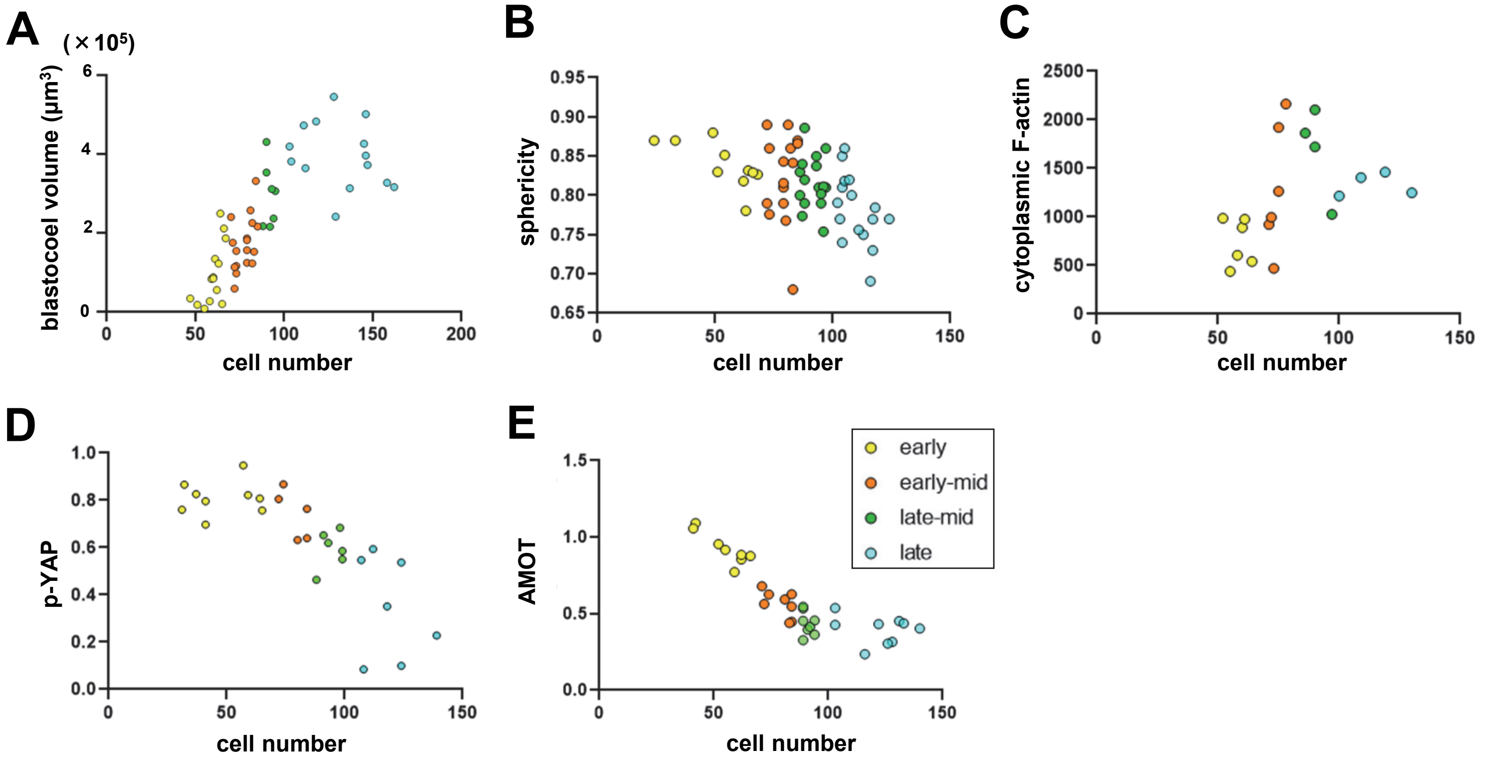

Figure S2

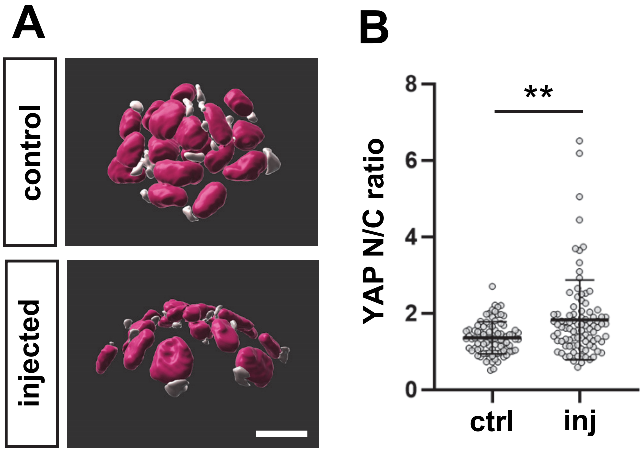

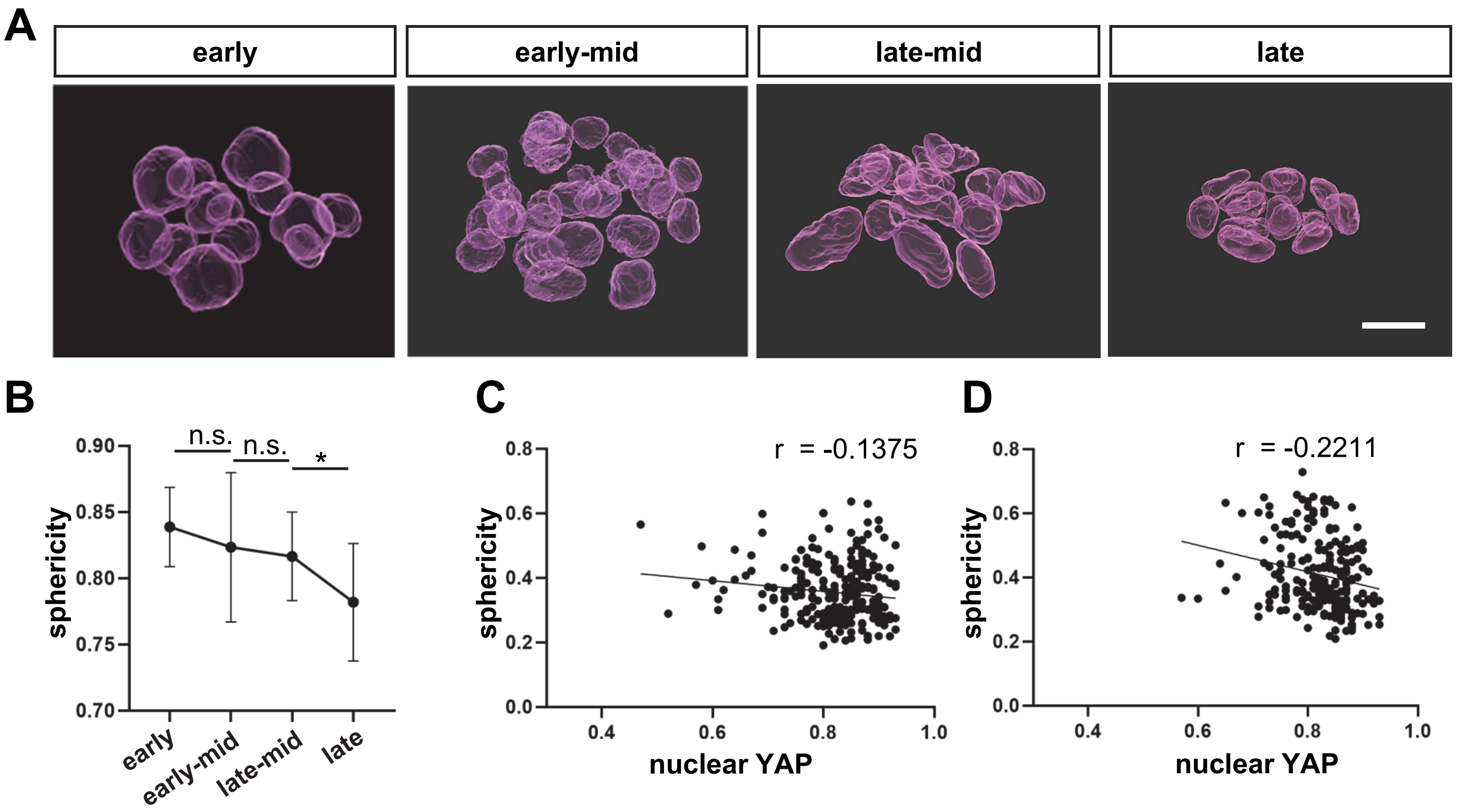

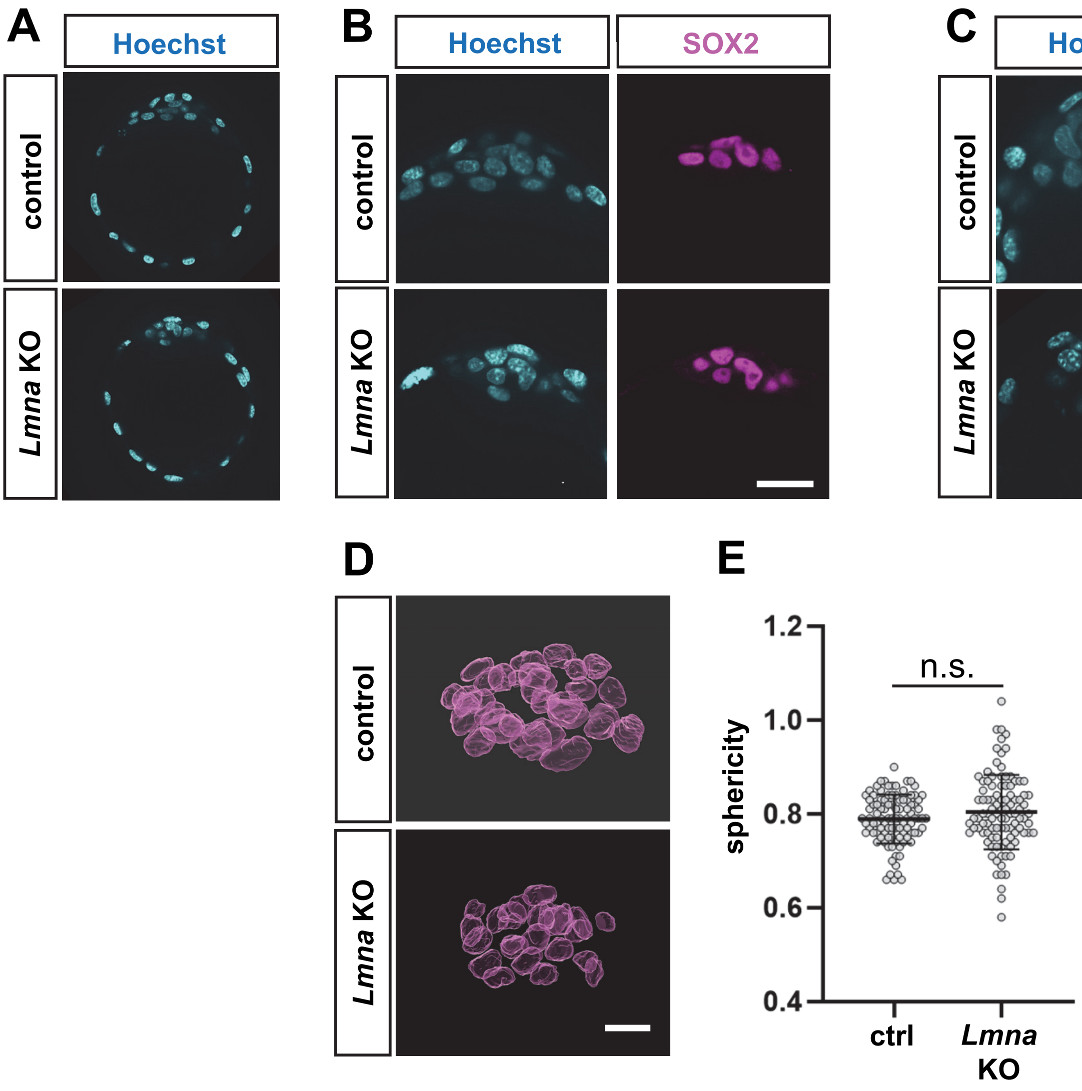

Figure S5

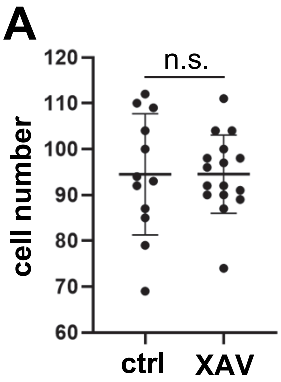

Figure S6

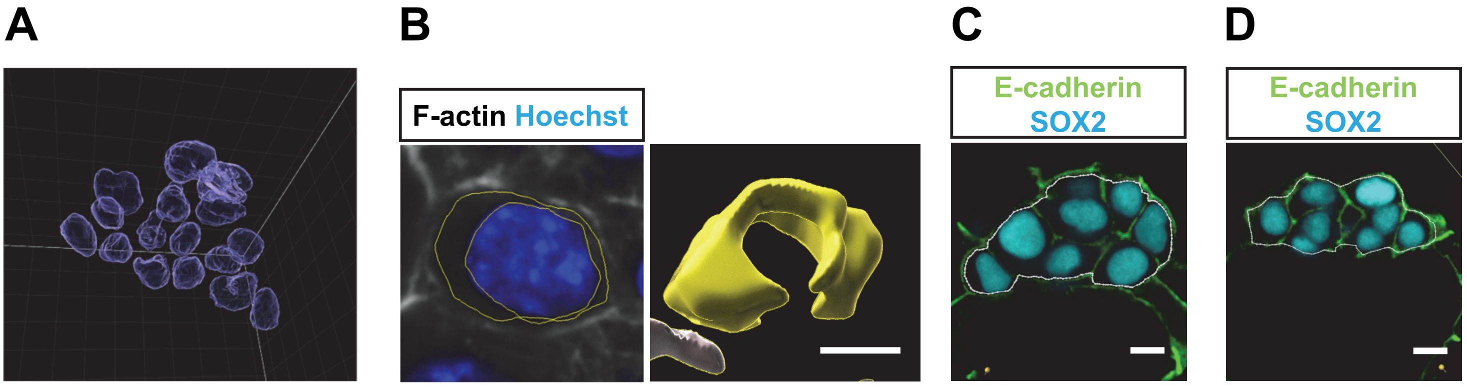
